# A CRISPR-based SARS-CoV-2 diagnostic assay that is robust against viral evolution and RNA editing

**DOI:** 10.1101/2020.07.03.185850

**Authors:** Kean Hean Ooi, Jie Wen Douglas Tay, Seok Yee Teo, Mengying Mandy Liu, Pornchai Kaewsapsak, Shengyang Jin, Yong-Gui Gao, Meng How Tan

## Abstract

Extensive testing is essential to break the transmission of the new coronavirus SARS-CoV-2, which causes the ongoing COVID-19 pandemic. Recently, CRISPR-based diagnostics have emerged as attractive alternatives to quantitative real-time PCR due to their faster turnaround time and their potential to be used in point-of-care testing scenarios. However, existing CRISPR-based assays for COVID-19 have not considered viral genome mutations and RNA editing in human cells. Here, we present the VaNGuard (Variant Nucleotide Guard) test that is not only specific and sensitive for SARS-CoV-2, but can also detect the virus when its genome or transcriptome has evolved or has been edited by deaminases in infected human cells. We show that an engineered AsCas12a enzyme is more tolerant of mismatches than wildtype LbCas12a and that multiplexed Cas12a targeting can overcome the presence of single nucleotide variations. Our assay can be completed in 30 minutes with a dipstick for a rapid point-of-care test.

## Introduction

COVID-19 is an ongoing global pandemic caused by SARS-CoV-2, a novel coronavirus of zoonotic origin. The outbreak was first reported in Wuhan, China^1-3^ and has since spread to more than 200 countries in all continents. As of 26 May 2020, there are over 5.4 million confirmed cases and 340,000 deaths worldwide, underscoring the severity of the disease. Importantly, given the high human-to-human transmission potential of SARS-CoV-2 including from asymptomatic carriers^4-6^, rapid and accurate diagnosis is critical for timely treatment and outbreak control. Currently, quantitative real-time PCR (qRT-PCR) is the gold standard method to detect COVID-19. However, it requires specialized and expensive instrumentation to run and thus must be carried out in dedicated facilities with the necessary equipment and expertise. Furthermore, the turnaround time for qRT-PCR is too slow. Even excluding the time it takes to transfer samples from collection points to the test facilities, the PCR process itself will take more than an hour to run. Besides qRT-PCR, serological tests that detect antibodies against SARS-CoV-2 are actively under development. However, such tests have limited practical use for identifying infectious individuals as antibodies are only detectable in later stages of infection when opportunities to treat and limit disease transmission have passed. Hence, there is still an unmet need for rapid, specific, and sensitive point-of-care tests (POCT) for SARS-CoV-2 that are easy to use and can be performed in resource-limited settings.

CRISPR-Cas has emerged as a powerful technology that can potentially drive next-generation diagnostic platforms. After binding to and cutting a specific target substrate, certain Cas enzymes are then hyperactivated to cleave all neighbouring nucleic acids indiscriminately^7-9^. By programming the Cas nuclease to recognize desired sequences, such as those containing cancer mutations or from pathogens-of-interest, and providing single-stranded DNA (ssDNA) or RNA reporter molecules in the reaction mix, various groups have successfully developed CRISPR-based diagnostics (CRISPR-Dx) for a range of applications^9- 14^. Unsurprisingly, it has also not escaped attention that the same technology can be quickly applied to tackle the ongoing COVID-19 outbreak. Within a few months, nine different CRISPR-based assays have been announced so far (Table 1)^15-23^, underscoring the ease-of-use and versatility of the technology.

**Table 1.** List of CRISPR-based assays for COVID-19 that have been announced (as of 26 May 2020).

| Name | DETECTR | CARMEN | CASdetec | STOPCovid | CRISPR-nCoV | CRISPR-Detection | AIOD-CRISPR | FELUDA | CREST |
| --- | --- | --- | --- | --- | --- | --- | --- | --- | --- |
| Status | Published<sup>15</sup> | Published<sup>16</sup> | Published<sup>17</sup> | MedRxiv preprint<sup>18</sup> | MedRxiv preprint<sup>19</sup> | BioRxiv preprint<sup>20</sup> | BioRxiv preprint<sup>21</sup> | BioRxiv preprint<sup>22</sup> | BioRxiv preprint<sup>23</sup> |
| Enzyme | LbCas12a | LwCas13a | AapCas12b | AapCas12b | LwCas13a | LbCas12a | LbCas12a | FnCas9 | LwCas13a |
| Based on collateral activity | Yes | Yes | Yes | Yes | Yes | Yes | Yes | No | Yes |
| Validated on patient samples | Yes (N = 11) | No | No | Yes (N = 12) | Yes (N = 52) | No | No | Yes (N = 1) | No |
| Affiliated company | Mammoth (partnering with GSK) | - | - | Sherlock | - | CASPR | - | Tata Sons | - |

While promising, all the existing CRISPR-Dx for COVID-19 have not considered the possibility that the viral sequences may be altered over time or in human cells. Viruses are known to mutate especially under selective pressure. Thousands of SARS-CoV-2 genomes had been sequenced and deposited in the GISAID database^24,25^ and analysis of their sequences revealed numerous mutations, suggesting an ongoing adaptation of the coronavirus to its novel human host^26^. In particular, mutations had been discovered in the target sites of many current COVID-19 diagnostic tests and could affect the performance of these qRT-PCR tests^27^. Moreover, mutations in the SARS-CoV-2 genome may also create mismatches in the guide RNA (gRNA) binding site and consequently affect the Cas ribonucleoprotein (RNP) complex’s ability to recognize its target. In addition, ADAR and APOBEC deaminases form part of the human host’s innate immune responses to viral infection and had recently been shown to edit SARS-CoV-2 RNA^28^. The respective adenosine-to-inosine and cytosine-to-uracil changes may also affect the ability of the CRISPR-Cas system to detect the virus.

Here, we report the development of CRISPR-Dx for COVID-19 that incorporate design features that mitigate the loss in signal caused by genomic mutations or RNA editing. We screened several different Cas12a enzymes and found that enAsCas12a, an engineered variant of AsCas12a^29^, was able to tolerate mismatches at the target site better than wildtype Cas12a nucleases. Furthermore, we demonstrated that incorporation of two gRNAs into the CRISPR-Cas system resulted in partial rescue of the output signal when a variant nucleotide was present in the substrate. Notably, while our assay could tolerate single nucleotide variations in the target sites, it still maintained high specificity and was able to distinguish SARS-CoV-2 from SARS-CoV and MERS-CoV reliably. Taken together, our VaNGuard (<u>Va</u>riant<u>N</u>ucleotide <u>Guard</u>) test holds the potential to address the need for a robust and rapid diagnostic assay that will help arrest viral spread and enable worldwide economies to re-open safely in the pandemic.

## Results

### Characterization of an existing N-gene gRNA

We started off by examining the design of the earliest CRISPR-based assay for COVID-19^15^. In this DETECTR assay, wildtype LbCas12a was paired with a 20-nucleotide (nt) gRNA targeting the N-gene of SARS-CoV-2. Given that various Cas12 enzymes had been successfully utilized for human genome engineering, we asked if LbCas12a was the best nuclease to deploy in a diagnostic test. To this end, we purified five different Cas12a enzymes and then paired each one of them with the same gRNA (herein termed N-Mam gRNA) in a fluorescence trans-cleavage assay (Fig. 1a). To assess the feasibility of a CRISPR-based diagnostic assay being deployed in a non-laboratory setting (e.g. a home setting), we carried out the reactions at room temperature using a 500-base pair (bp) synthetic DNA fragment of the SARS-CoV-2 N gene. Fluorescence was monitored over the course of 30 minutes in a microplate reader (Supplementary Fig. 1). Expectedly, LbCas12a was able to clearly detect SARS-CoV-2 with no cross-reactivity for SARS-CoV or MERS-CoV at the end of the reaction. Nevertheless, the other four Cas12a enzymes also performed similarly to LbCas12a, with enAsCas12a yielding an even higher fluorescence signal than LbCas12a in the presence of the intended SARS-CoV-2 substrate (Fig. 1b). We also found that the minimum spacer length for the N-Mam gRNA was 20nt, as shortening of the spacer led to a reduction in fluorescence output for all the tested nucleases.

**Fig. 1.**
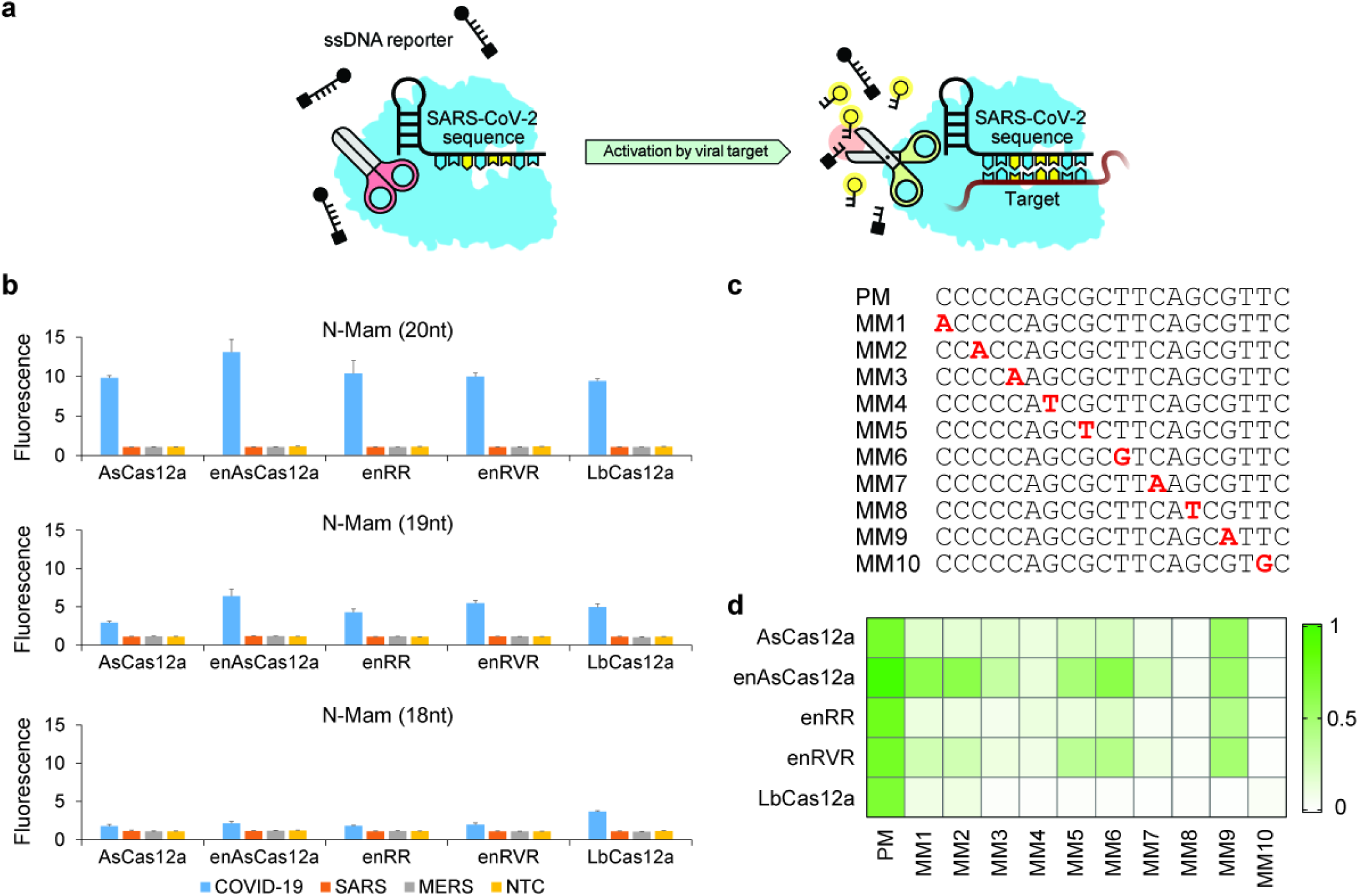
Activity and mismatch tolerance of various Cas12a enzymes with the N-Mam gRNA. **a** Schematic of a fluorescence trans-cleavage assay. Here, the reporter comprises a fluorophore linked to a quencher by a short piece of ssDNA. The gRNA is programmed to recognize a particular locus of the SARS-CoV-2 genome. In the absence of the virus, the reporter molecule is intact and thus no fluorescence is observed. However, when the virus is present, the Cas12a RNP will bind to and cleave its programmed target, become hyperactivated, and cut the ssDNA linker between the fluorophore and quencher, thereby generating a fluorescence signal. **b** Fluorescence measurements using a microplate reader after 30 minutes of cleavage reaction. 1E11 copies of DNA template corresponding to one of the coronaviruses (see colour bar) were present in a 50μl reaction. Spacers of three different lengths targeting the N-Mam locus were tested. There was no cross-reactivity for SARS-CoV or MERS-CoV as expected. All the measurements were normalized to the no-template control (NTC) at the start of the experiment. Data represent mean ± s.e.m. (n = 3 biological replicates). **c** Sequences of perfect matched (PM) or mismatched (MM) spacers targeting the N-Mam locus. Each mismatched position is indicated by a bold red letter. **d** Heatmap showing the tolerance of various Cas12a enzymes to mismatched N-Mam gRNAs. The fluorescence readings are scaled between 0 and 1, where 1 is the highest measurement obtained and 0 is the background signal for NTC at the start of the experiment.

Next, we tested how mismatches at the gRNA-substrate interface may affect the fluorescence signal. We generated ten new gRNAs targeting the same N-Mam locus, with each harbouring a single point mutation at variable locations along the spacer (Fig. 1c). From the trans-cleavage assay, we observed that LbCas12a was very sensitive to imperfect base pairing between the gRNA and the substrate, as any mismatch along the spacer reduced the fluorescence output to near-background levels (Fig. 1d and Supplementary Fig. 2). In contrast, AsCas12a and its variants were able to tolerate a mismatch 4nt from the 3’ end of the gRNA (MM9). Furthermore, we found that enAsCas12a was the most tolerant to mismatches among the five tested enzymes. Hence, our results suggest that enAsCas12a should be paired with the N-Mam gRNA to safeguard against viral mutations or RNA editing at the target site.

**Fig. 2.**
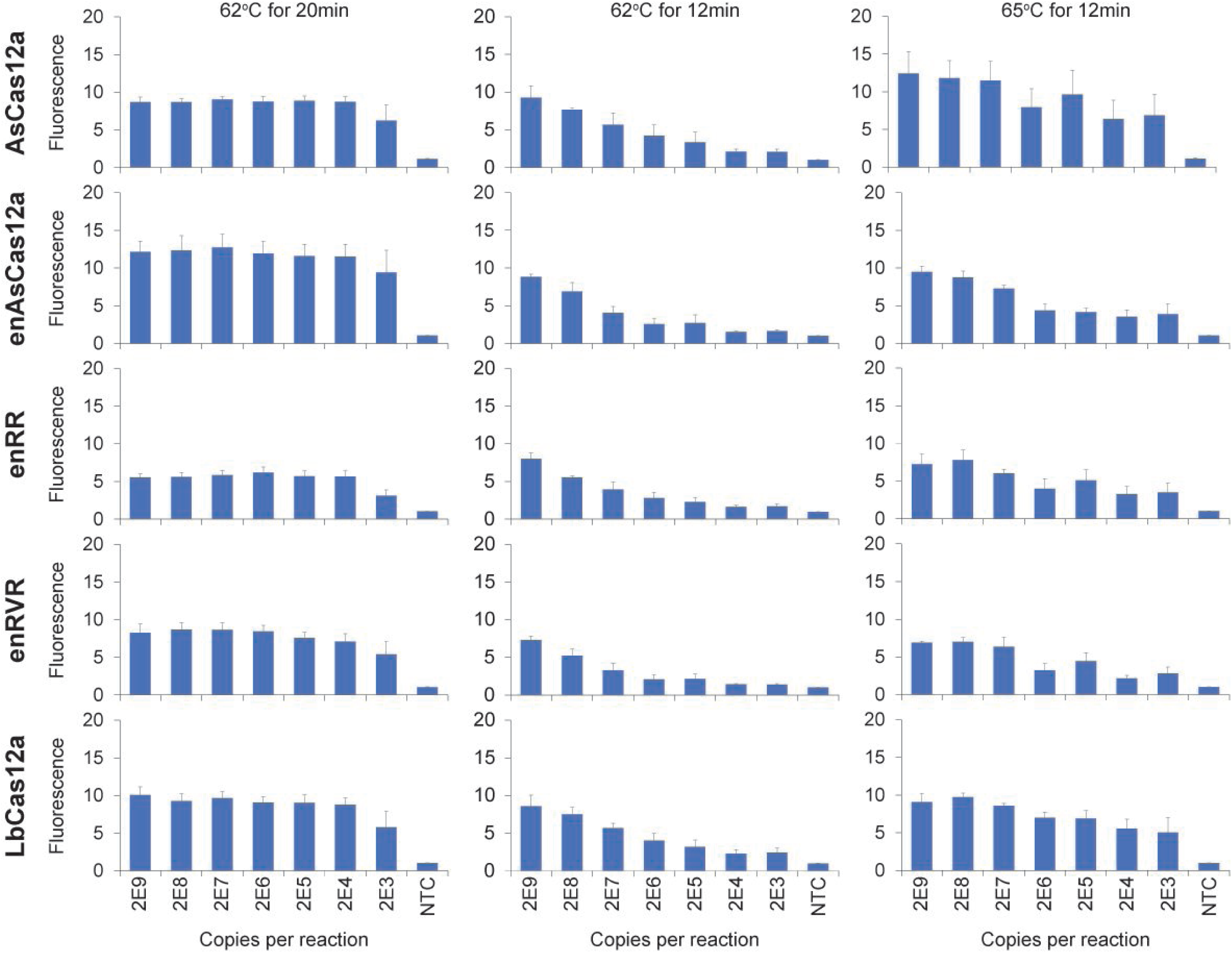
Analytic limit of detection (LoD) under various LAMP reaction conditions. Different copies of synthetic SARS-CoV-2 RNA fragments were used as input and the volume of each RT-LAMP reaction was 25μl. The fluorescence readings here were taken after 10 minutes of cleavage reaction. Data represent mean ± s.e.m. (62°C for 20min: n = 4-7 biological replicates; 62°C/ 65°C for 12min: n = 3 biological replicates).

### Evaluation of RT-LAMP parameters

To enhance sensitivity, CRISPR-Cas detection is typically combined with an isothermal amplification step, of which there are several options. Due to supply chain issues in the ongoing pandemic, reverse transcription loop-mediated isothermal amplification (RT-LAMP)^30^ is the method-of-choice for COVID-19 applications. In the DETECTR assay, the RT–LAMP reaction was performed at 62°C for 20–30 minutes^15^. Therefore, we first carried out RT-LAMP at 62°C for 20 minutes on synthetic *in vitro*-transcribed (IVT) SARS-CoV-2 RNA templates and then used the amplified products in our fluorescence trans-cleavage assay (Fig. 2 and Supplementary Fig. 3). For all the tested enzymes, the viral sequence was consistently and clearly detected in every replicate when 20,000 or more copies of RNA were used as input to RT-LAMP. However, in contrast to the published report^15^, when 2,000 copies of RNA were used in a 25μl RT-LAMP reaction (i.e. 80 copies per μl) instead, the viral sequence was detected in only around half the replicates for all the Cas enzymes, including LbCas12a.

**Fig. 3.**
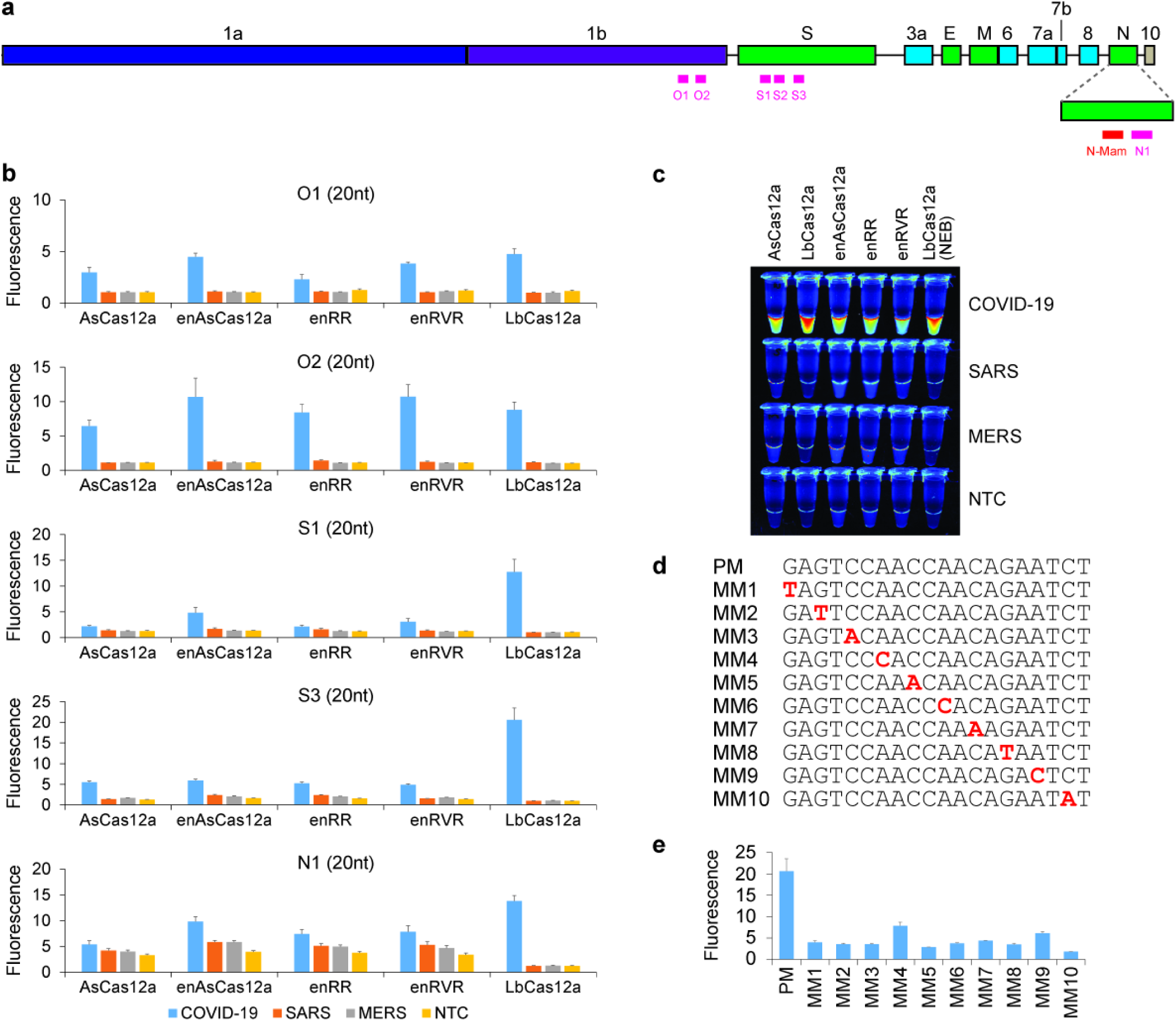
Collateral activity of various Cas12a enzymes complexed with different gRNAs. **a** Organization of the SARS-CoV-2 genome. ORF1a and ORF1b occupy over half of the genome. Genes encoding structural proteins are indicated by green boxes, while genes encoding accessory proteins are indicated by cyan boxes. Although ORF10 is annotated in the genome, there is currently no evidence of its expression^50^. The locations of the new gRNAs are shown by pink bars below the genes, while the N-Mam locus is shown by a red bar. **b** Fluorescence measurements using a microplate reader after 30 minutes of cleavage reaction. 1E11 copies of the relevant DNA target were present in a 50μl reaction. All the readings were normalized to the negative control (NTC) at the start of the experiment. The N1 gRNA gave an unexpected result, whereby it triggered the collateral activity of AsCas12a and its variants even in the absence of a template. Data represent mean ± s.e.m. (n = 3-6 biological replicates). **c** Visualization of cleavage reactions for the S3 gRNA using a UV transilluminator. Transition of colours from blue to yellow to red indicates an increasing amount of collateral activity. **d** Sequences of perfect matched (PM) or mismatched (MM) spacers targeting the S3 locus. Each mismatched position is indicated by a bold red letter. **e** Collateral activity of LbCas12a complexed with perfect matched or mismatched S3 gRNA. The fluorescence measurements were taken after 30 minutes of cleavage reaction using a microplate reader and all the readings were normalized to the NTC at the start of the experiment. Our results indicate that LbCas12a should be paired with the S3 gRNA instead of the N-Mam gRNA if a diagnostic assay that is highly specific for known SARS-CoV-2 isolates is desired. Data represent mean ± s.e.m. (n = 3 biological replicates).

Subsequently, we asked how the sensitivity of the CRISPR-based assay would be affected if we varied the parameters of the LAMP reaction (Fig. 2 and Supplementary Fig. 3). When we decreased the duration of LAMP from 20 to 12 minutes, we observed that the fluorescence signal showed an exponential decay with every 10-fold reduction in RNA copy number, indicating that the analytic limit of detection (LoD) became poorer if the amplification step was performed for too short a period of time. Interestingly however, when we increased the reaction temperature from 62°C to 65°C while keeping the duration at 12 minutes, we were able to partially recover the fluorescence signal. Further increase in temperature from 65°C to 68°C caused a deterioration in the performance of our CRISPR-Dx (Supplementary Fig. 4). Hence, our results suggest that 65°C is the optimal temperature for LAMP and that the amplification reaction is sensitive to small variations in temperature.

**Fig. 4.**
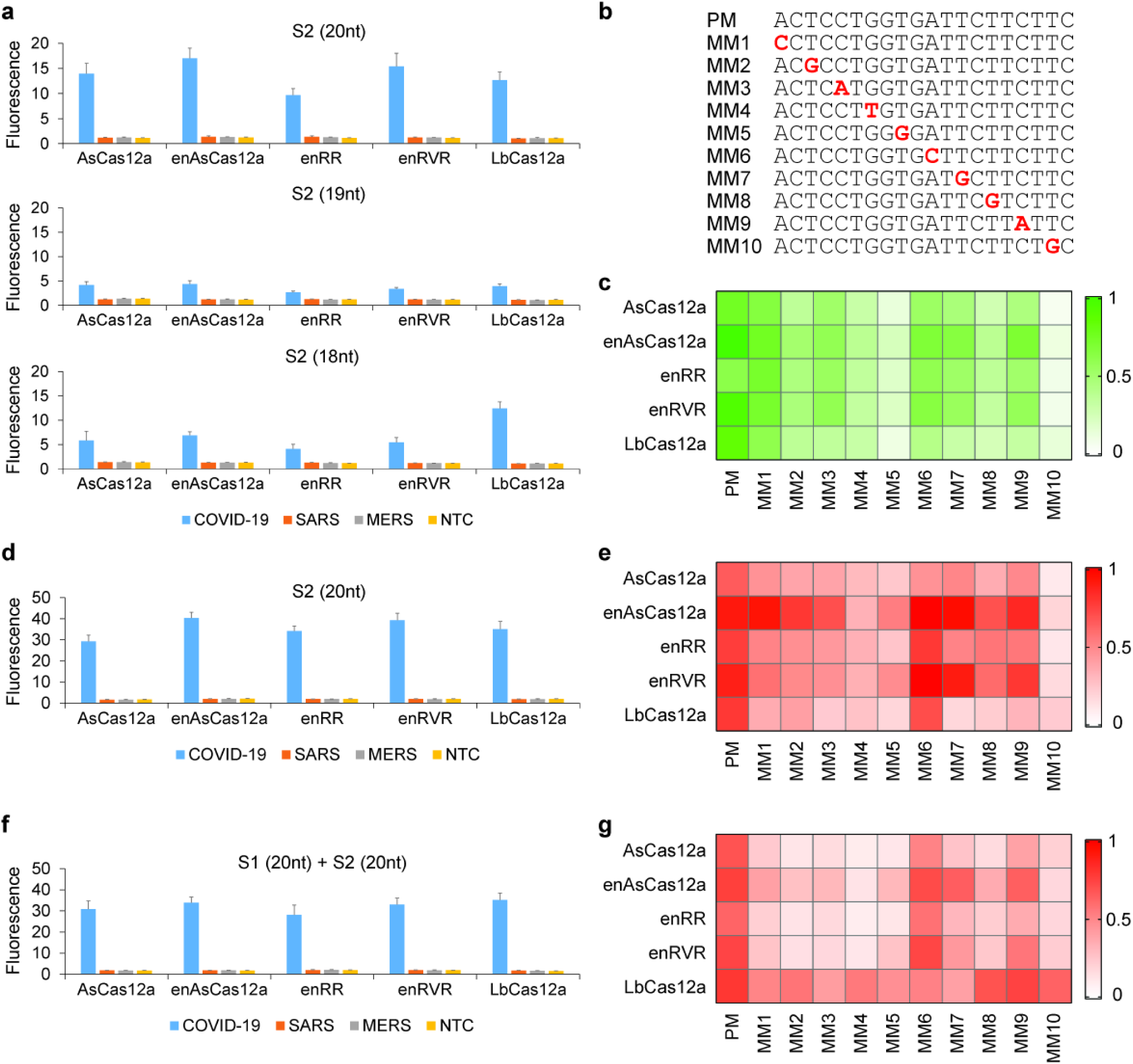
Activity and mismatch tolerance of various Cas12a enzymes with the S2 gRNA. **a** Fluorescence measurements for a single S2 gRNA after 30 minutes of trans-cleavage reaction at 24°C. 1E11 copies of DNA template corresponding to one of the three coronaviruses (see colour bar) were present in a 50μl reaction. Spacers of three different lengths targeting the S2 locus were tested. All the measurements were normalized to the negative control (NTC) at the 0min timepoint. No cross-reactivity for SARS-CoV or MERS-CoV was detected. Unexpectedly, the 18nt spacer yielded higher fluorescence than the 19nt spacer for all the tested nucleases, especially LbCas12a. Data represent mean ± s.e.m. (n = 3-4 biological replicates). **b** Sequences of perfect matched (PM) or mismatched (MM) spacers targeting the S2 locus. Each mismatched position is indicated by a bold red letter. **c** Heatmap showing the tolerance of various Cas12a enzymes to mismatched S2 gRNAs when the trans-cleavage assay was performed at 24°C. The fluorescence readings are scaled between 0 and 1 (variable shades of green), where 1 is the highest measurement obtained at 24°C and 0 is the background signal for NTC at the start of the experiment. **d** Fluorescence measurements for a single S2 gRNA after 30 minutes of trans-cleavage reaction at 37°C. There was still no cross-reactivity for SARS-CoV or MERS-CoV at the higher temperature, but the fluorescence signal for SARS-CoV-2 was approximately twice as high. Data represent mean ± s.e.m. (n = 3 biological replicates). **e** Heatmap showing the tolerance of various Cas12a enzymes to mismatched S2 gRNAs when the trans-cleavage assay was performed at 37°C. The fluorescence readings are scaled between 0 and 1 (variable shades of red), where 1 is the highest measurement obtained at 37°C and 0 is the background signal for NTC at the start of the experiment. The most detrimental mismatch position appears to be 2nt from the PAM-distal end (MM10). **f** Fluorescence measurements for two PM gRNAs (S1 and S2) after 30 minutes of cleavage reaction at 37°C. No cross-reactivity for SARS-CoV or MERS-CoV was observed. Data represent mean ± s.e.m. (n = 3 biological replicates). **g** Heatmap showing how the addition of a second perfect matched S1 gRNA changed the tolerance of various Cas12a enzymes to mismatched S2 gRNAs. The trans-cleavage assay was performed at 37°C, with the fluorescence readings scaled between 0 and 1.

### Screening of additional gRNAs

We wondered if there may be other gRNAs that could yield a stronger fluorescence signal than the N-Mam gRNA in our trans-cleavage assay. The genome of SARS-CoV-2 contains two open reading frames (ORFs), ORF1a and ORF1b, that encode multiple non-structural proteins (nsps), four genes that encode conserved structural proteins (spike protein [S], envelope protein [E], membrane protein [M], and nucleocapsid protein [N]), and several accessory ORFs of unclear function (Fig. 3a)^31^. We aligned the genomes of SARS-CoV-2, SARS-CoV, and MERS-CoV and selected six additional target sites (O1, O2, S1, S2, S3, and N1) that not only contained the necessary TTTV protospacer adjacent motif (PAM) for Cas12a but were also highly divergent between the three coronaviruses (Supplementary Fig. 5). We then analysed the collateral activities of our five Cas12a enzymes paired with each of the six new gRNAs over the course of 30 minutes using synthetic DNA fragments of the relevant genes as targets. Curiously, when the N1 gRNA was paired with either wildtype AsCas12a or any of its engineered variants (enAsCas12a, enRR, and enRVR)^29^, we were able to detect some fluorescence signal even in the absence of template (Fig. 3b and Supplementary Fig. 6), suggesting that AsCas12a may be hyper-activated by certain gRNAs *in vitro* without the need for it to cleave its intended target. Such a phenomenon was not observed with LbCas12a.

**Fig. 5.**
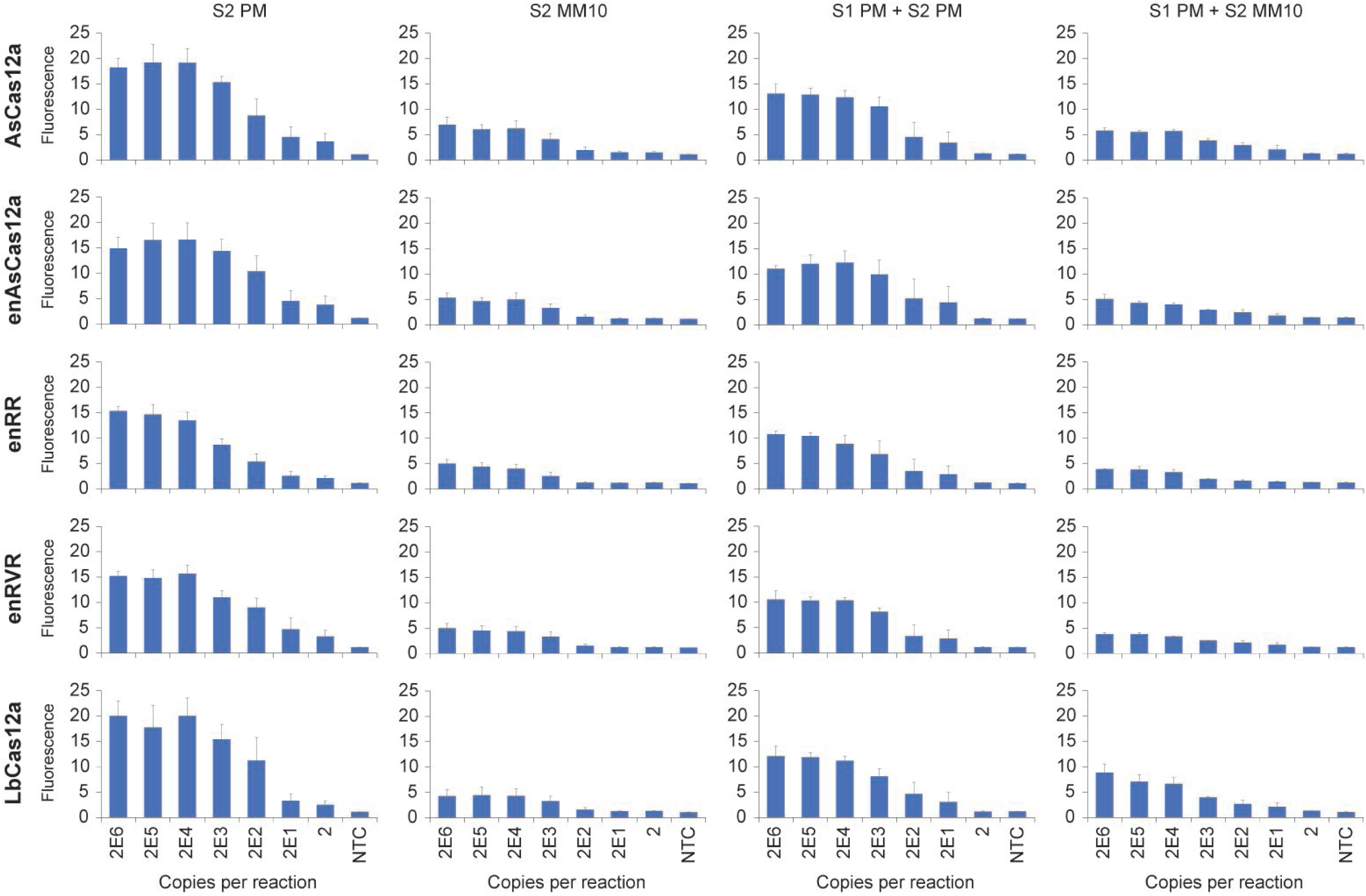
Analytic LoD for gRNAs targeting the S-gene. Different copies of *in vitro* transcribed SARS-CoV-2 RNA fragments were used as input to the RT-LAMP reaction, which was performed at 65°C for 15 minutes. The Cas detection reaction was then carried out at 37°C, with the fluorescence measurements here taken after 10 minutes using a microplate reader. Data represent mean ± s.e.m. (n = 3-5 biological replicates).

**Fig. 6.**
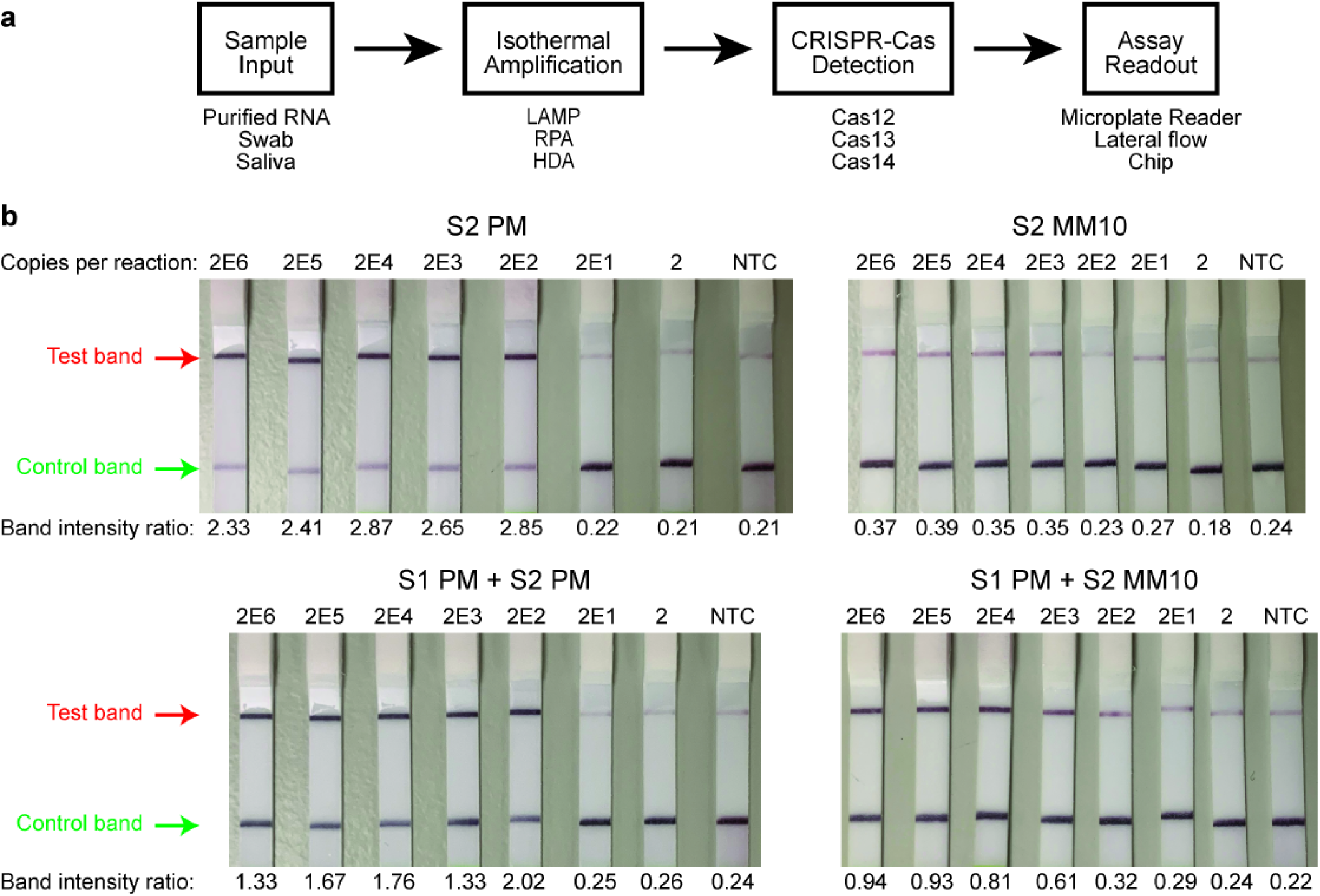
Implementation of our VaNGuard assay on lateral flow strips. **a** Overview of a prototypical CRISPR-Dx workflow. While a microplate reader can allow up to 96 samples to be processed at once, it is not amenable to point-of-care testing. In contrast, a lateral flow strip proves a simple visual readout akin to an off-the-shelf pregnancy test. **b** Detection of SARS-CoV-2 sequence using gRNAs targeting the S-gene. Different copies of synthetic SARS-CoV-2 RNA fragments were used as input to the RT-LAMP reaction, which was performed at 65°C for 15 minutes. Next, the Cas detection reaction was carried out at 37°C for 10 minutes before a dipstick was added to each reaction tube. The bands on the dipstick appeared by 2 minutes. In total, the VaNGuard assay was completed in under 30 minutes.

Overall, the three S-gene gRNAs were able to generate similar or higher fluorescence signals than the N-Mam gRNA with the appropriate Cas12a nuclease, while the two O-gene gRNAs generally performed worse. In particular, in the presence of the SARS-CoV-2 target, the collateral activity of LbCas12a complexed with the S3 gRNA was approximately double that of the same enzyme complexed with the N-Mam gRNA (Fig. 3b and Supplementary Fig. 6). There was no cross-reactivity for SARS-CoV or MERS-CoV. We also confirmed that the activity of our own purified protein was comparable to that of a commercially available LbCas12a enzyme used in several recently announced CRISPR-based COVID-19 diagnostic assays^15,20,21^ (Fig. 3c). Subsequently, we investigated the mismatch tolerance of LbCas12a at the S3 target locus but found that collateral activity was greatly diminished for all the mismatched (MM) gRNAs tested (Fig. 3d, e and Supplementary Fig. 7). Hence, although the S3 gRNA may be used in a highly specific assay for known SARS-CoV-2 isolates, it is not ideal for a diagnostic assay that is robust against potential mutations at the target site.

### Detailed characterization of a new S-gene gRNA

Next, we turned our attention to the S2 gRNA. In the presence of the SARS-CoV-2 substrate, this gRNA generated stronger fluorescence signals than the N-Mam gRNA when paired with LbCas12a, AsCas12a, enAsCas12a, or enRVR (Fig. 4a and Supplementary Fig 8). The engineered enAsCas12a enzyme exhibited the highest collateral activity with the S2 gRNA. Importantly, no cross-reactivity for SARS-CoV or MERS-CoV was observed regardless of the Cas12a nuclease used. Furthermore, we found that the minimum spacer length for the S2 gRNA was 20nt, as shortening of the spacer reduced the collateral activity of all tested nucleases, although unexpectedly, the 18nt gRNA gave higher fluorescence signals than the 19nt gRNA.

We sought to determine how mismatches at the S2 gRNA-substrate interface may affect the collateral activity of the Cas12a enzymes. To this end, we generated ten additional gRNAs with each harbouring a point mutation at different positions along the spacer (Fig. 4b). Interestingly, we found from the trans-cleavage assay that individual mismatches at the S2 locus affected the collateral activity of all the Cas12a endonucleases much less than those at the N-Mam locus (Fig. 4c and Supplementary Fig. 9). Nevertheless, enAsCas12a and enRVR again exhibited the highest tolerance for mismatches, while wildtype LbCas12a was again the most sensitive to imperfect base pairing between the gRNA and its target substrate.

So far, we had performed the CRISPR-Cas detection at room temperature (24°C) to simulate a non-laboratory setting, but we wondered how the performance of our diagnostic assay would improve if the detection was performed at the optimal temperature (37°C) of the Cas12a enzymes instead. From the trans-cleavage assay, we observed that the fluorescence signal increased approximately twice as fast at 37°C for all tested enzymes in the presence of the intended SARS-CoV-2 template and reached significantly higher levels after 30 minutes of reaction (P < 0.01, Student’s t-test) (Fig. 4d and Supplementary Fig. 10). There was again no cross-reactivity for SARS-CoV and MERS-CoV. In addition, the activity profile in the presence of point mutations remained similar and in fact even improved slightly for all the nucleases with respect to mismatch tolerance (Fig. 4e and Supplementary Fig. 10). Taken together, our results indicate that our CRISPR-based assay should be performed at 37°C if a faster test result is desired and also suggest that the S2 gRNA is more suitable than the N-Mam gRNA in a diagnostic assay that guards against viral evolution or intracellular RNA editing.

### Multiplex Cas12a targeting

Besides searching for Cas-gRNA pairs that are not only specific for SARS-CoV-2 but are also tolerant of mismatches at the binding site, another strategy to enhance robustness is to incorporate two or more distinct gRNAs into the detection module. As a proof-of-concept, we sought to buffer the collateral activity of LbCas12a, which appeared to be more sensitive to imperfect base pairing at the gRNA-target interface than AsCas12a and its engineered variants. Moreover, LbCas12a protein is commercially available and can be readily bought by diagnostic laboratories that do not have the ability to purify their own enzymes. To keep all the target sites within the same LAMP products, we may pair the S2 gRNA with either the S1 or the S3 gRNA, as both worked well with LbCas12a (Fig. 3b and Supplementary Fig. 6). We then tried to design LAMP primers using the online software PrimerExplorer V5. However, we were unable to find any set of primers that would produce an amplicon smaller than 500bp that contained both the S2 and S3 loci. In contrast, many primer sets could be obtained for S1 and S2. Hence, we decided to perform our multiplexed targeting experiments with the S1 and S2 gRNAs.

Using a synthetic DNA fragment of the SARS-CoV-2 S gene as substrate, we first assessed the collateral activity of our five purified Cas12a enzymes when they were combined with both the S1 and S2 gRNAs. All five nucleases exhibited robust activity with perfect matched (PM) gRNAs (Fig. 4f and Supplementary Fig. 11). However, we noted that addition of the S1 gRNA caused a small but obvious reduction in the trans-cleavage activity of the three AsCas12a variants, while that of LbCas12a remained approximately the same. This was likely because the S1 gRNA would compete with the S2 gRNA for the Cas proteins but only LbCas12a exhibited strong activity with the S1 gRNA (Fig. 3b and Supplementary Fig. 6). Subsequently, we profiled the activity of the Cas12a nucleases when the S1 PM gRNA was used together with each one of the S2 MM gRNAs (Fig. 4g and Supplementary Fig. 11). Strikingly, we now observed that LbCas12a exhibited the best overall tolerance for mismatches, while AsCas12a and its engineered variants became more sensitive to the mismatches.

Next, we sought to determine the analytic limit of detection (LoD) of our CRISPR-Dx. We tested three sets of LAMP primers and found one set that amplified well even with low amounts of template (Supplementary Fig. 12). Therefore, we carried out RT-LAMP on synthetic *in vitro*-transcribed (IVT) SARS-CoV-2 RNA templates using this selected primer set. The reaction was performed at 65°C for 15 minutes because we had earlier found the temperature to be optimal for LAMP and we sought to minimize the duration of our diagnostic test. The amplified products were used immediately in our trans-cleavage assay, with the fluorescence monitored over time in a microplate reader (Fig. 5 and Supplementary Fig. 13). When only a single S2 PM gRNA was used, we found that the LoD was around 200 copies per reaction for all tested Cas12a nucleases. Expectedly, introduction of a single mismatch (MM10) at the gRNA-substrate interface worsened the sensitivity of CRISPR-Cas detection. More importantly, addition of a S1 PM gRNA was able to partially rescue the mismatch at the S2 locus for LbCas12a, but not for the other Cas12a enzymes. Taken together, our trans-cleavage assays revealed that the use of two gRNAs could increase the robustness of CRISPR-Dx with respect to the presence of variant nucleotides, but care must be taken to only utilize gRNAs that would work well together with the selected Cas nuclease.

### Towards a point-of-care test

To broaden the use cases of our diagnostic assay, we sought to develop a portable point-of-care test (POCT). After the CRISPR-Cas trans-cleavage reaction has taken place, the results can be read out in different ways (Fig. 6a). While a microplate reader is useful for high throughput screening of samples in a centralized facility, it is not amenable to non-laboratory settings like ports of entry, workplaces, schools, public spaces, and homes. Hence, we decided to visualize the results of our assay on a lateral flow strip (Supplementary Fig. 14). Here, the reporter molecule consists of a fluorescent dye (fluorescein) linked to biotin by a short piece of ssDNA. An anti-fluorescein antibody conjugated to gold binds to the dye on the strip. When the viral substrate is absent, the reporter is intact and captured by streptavidin at the control line. However, when the viral target is present, the reporter is cleaved and the fluorescein-antibody complex migrates to the test line where it is captured by an immobilized secondary antibody.

We performed the RT-LAMP reaction at 65°C for 15 minutes followed by the CRISPR-Cas trans-cleavage reaction at 37°C for 10 minutes before adding a lateral flow strip to the reaction tube. Bands appeared at either the test line or the control line within two minutes (Fig. 6b). Hence, in total, the entire assay took slightly under 30 minutes to complete. Here, we focused on LbCas12a to demonstrate the utility of multiplex targeting. The S2 gRNA (PM or MM10) was used in the assay with or without a second S1 PM gRNA. We also tested different copy numbers of the synthetic SARS-CoV-2 RNA template. Overall, we found that our lateral flow assays gave results that mirrored those from a microplate reader. When the S2 PM gRNA was used alone or in conjunction with the S1 PM gRNA, dark bands appeared at the test line in the presence of at least 200 copies of template in the reaction mix. Expectedly, without any multiplexing, introduction of a mismatch at the S2 gRNA-target interface dramatically reduced the intensity of the test bands and worsened the analytic LoD to 2,000 copies per reaction. Importantly, addition of the S1 PM gRNA was able to partially restore the intensity of the test bands. Quantification of the test and control band intensities further provided a more objective measure of whether the intended target was detected or not. In the case of a mismatch at the S2 locus but perfect base pairing at the S1 locus, we were able to tell that 200 copies per reaction gave a positive test result based on the ratio of the test to control band intensities. In the future, a web or mobile phone application can be developed to distinguish such borderline cases.

## Discussion

There is an urgent healthcare need for rapid and accurate diagnostic tests for COVID-19. What initially started as an infectious disease confined to Wuhan, China has since spread to more than 200 countries worldwide within the span of a few months because SARS-CoV-2 has a high potential for human-to-human transmission and the majority of carriers are asymptomatic or only mildly ill. At present, qRT-PCR serves as the gold standard for viral detection, with dozens of test kits available in the market. However, the method requires dedicated instrumentation and trained operators and also has a slow turnaround time. Hence, there is still an unmet need for a rapid, sensitive, specific, and affordable SARS-CoV-2 diagnostic assay, which is essential for stopping viral spread and for the safe reopening of economies and schools.

CRISPR-Dx has the potential to meet society’s need for such a diagnostic test. The entire workflow consists of four main modules (Fig. 6a). First is the sample input. Although purified RNA is ideal for performance, the process of RNA extraction will take up precious time, increase cost, and stress the supply chain. Therefore, there is great interest in developing assays that can directly handle patient samples, including nasopharyngeal swabs and saliva. The second module is the isothermal amplification step, which is commonly implemented to enhance the sensitivity of CRISPR-Dx. LAMP^30^ is the method-of-choice in the current pandemic climate, as its reagents are readily available from several suppliers, but other approaches can also be used, including recombinase polymerase amplification (RPA)^32^ and helicase-dependent amplification (HDA)^33^. The third module is the CRISPR-Cas detection system. Most CRISPR-Dx rely on an indiscriminate collateral activity possessed by some Cas nucleases, including Cas12, Cas13, and Cas14 family members. Lastly, the fourth module is the assay readout. While our work here has demonstrated the use of a microplate reader (for high-throughput testing) and a lateral flow strip (for POCT), another possibility is a graphene-based field-effect transistor, whose high sensitivity has been reported to obviate the need for a pre-amplification step^34^. Overall, the cost of running a CRISPR-based test per sample is under S$9 (Table 2), which is around US$6.40 or €5.80 and is similar to that of an off-the-shelf pregnancy test. The bulk of the cost comes from the LAMP mastermix and the dipstick.

**Table 2.** Estimated cost of CRISPR-Dx for each sample.

| Reagent | Approximate Price (S\$) | Number of Reactions | Price per reaction (S\$) | |
| --- | --- | --- | --- | --- |
|  |  |  | Plate reader | Dipstick |
| RNA extraction kit | 1,200 | 90,800 | 0.01 | 0.01 |
| IVT Kit | 491 | 22,700 | 0.02 | 0.02 |
| RT-LAMP mastermix | 1,489 | 500 | 2.98 | 2.98 |
| Cleavage assay buffer | 41 | 1,000 | 0.04 | 0.04 |
| LbCas12a | 450 | 600 | 0.75 | 0.75 |
| LAMP primer (F3) | 5 | 1,200 | <0.01 | <0.01 |
| LAMP primer (B3) | 5 | 1,200 | <0.01 | <0.01 |
| LAMP primer (FIP) | 10 | 125 | 0.08 | 0.08 |
| LAMP primer (BIP) | 10 | 125 | 0.08 | 0.08 |
| LAMP primer (LF) | 5 | 300 | 0.02 | 0.02 |
| LAMP primer (LB) | 5 | 300 | 0.02 | 0.02 |
| FITC-quencher reporter | 315 | 650 | 0.48 | - |
| 96-well plate | 8 | 96 | 0.08 | - |
| FITC-biotin reporter | 200 | 5,555 | - | 0.04 |
| Dipstick | 490 | 100 | - | 4.90 |
| **TOTAL** |  |  | 4.58 | 8.94 |

While promising, current CRISPR-Dx assays for COVID-19 (Table 1) have not taken into account viral evolution and RNA editing. Alarmingly, mutations in the SARS-CoV-2 genome have been observed at the target sites of multiple existing qRT-PCR diagnostic tests^27^. Moreover, RNA editing mediated by the ADAR and APOBEC enzymes can also impact upon the performance of COVID-19 diagnostics. Hence, in this work, we sought to bolster the robustness of CRISPR-Dx against unexpected variant nucleotides introduced by evolutionary pressures or RNA editing. Starting from the DETECTR platform^9,15^, we tested several different natural and engineered Cas12a enzymes and found that enAsCas12a exhibited the highest tolerance for single mismatches at the gRNA-target interface. Importantly, high specificity for SARS-CoV-2 was still maintained with enAsCas12a, as no cross-reactivity for two other closely related coronaviruses SARS-CoV and MERS-CoV was observed. We also screened additional gRNAs and discovered that all the tested nucleases, except enRR, exhibited higher trans-cleavage activity with the S2 gRNA than with the N-Mam gRNA. Hence, our results indicate that enCas12a complexed with the S2 gRNA will serve as a more sensitive and robust SARS-CoV-2 detection system than the published LbCas12a and N-Mam gRNA pair. Nevertheless, we are mindful that enCas12a is not yet a commercially available enzyme. Hence, we demonstrated that a multiplex targeting strategy could also be utilized to enhance the robustness of CRISPR-Dx. For example, the LbCas12a nuclease, which is readily bought, may be combined with both the S1 and S2 gRNAs to increase the robustness of viral detection.

We note that diagnostic assays can be constructed out of isothermal amplification methods alone without coupling them to a separate CRISPR-Cas detection module. Such assays typically rely on the use of a turbidimeter to measure the extent of magnesium precipitation, labelled primers, or special dyes that sense pH changes, react with amplification by-products, or bind to double-stranded DNA (dsDNA). Due to their relative simplicity, over a dozen LAMP-only diagnostic assays for COVID-19 have been developed so far^35-49^. However, isothermal amplification frequently produces spurious non-specific products, which can give rise to false positive results. Although this problem may be circumvented by the use of a sequence-specific detection probe that is distinct from the LAMP primers^43^, the probe itself is not involved in any amplification process. In contrast, CRISPR-Dx confers two distinct rounds of specificity. The first round comes from primer-specific isothermal amplification such as LAMP, while the second round comes from gRNA-specific Cas detection. Furthermore, the CRISPR-Cas detection system is also capable of signal amplification because each hyperactivated Cas nuclease can proceed to cleave numerous reporter molecules. Hence, CRISPR-Dx can function like a photomultiplier tube and the assay duration can potentially be shortened if all the reagents are in a single-pot and the conditions are optimal for every reaction.

In conclusion, CRISPR-Dx can serve as a rapid, specific, sensitive, and affordable approach for the detection of SARS-CoV-2. Our work here has further provided two different strategies, namely the use of enCas12a and multiplex targeting, to enhance the robustness of the assay. It can be implemented in a high-throughput format through the use of a microplate reader or deployed as a POCT through the use of a lateral flow strip to enable us to halt viral transmission and reopen our society safely.

## Methods

### Plasmids and oligonucleotides

The pET28b-T7-Cas12a-NLS-6xHis expression plasmids were gifts from Keith Joung and Benjamin Kleinstiver (Addgene plasmid #114069 [AsCas12a], #114070 [LbCas12a], #114072 [enAsCas12a], #114075 [enRVR], and #114077 [enRR])^29^. DNA oligonucleotides, custom reporters for the trans-cleavage assays, and gene fragments (ORF1AB, S, and N) for the three coronaviruses SARS-CoV-2, SARS-CoV and MERS-CoV were synthesised by Integrated DNA Technologies. All oligonucleotides used in this study are listed in Supplementary File 1.

### Cas12a expression and purification

The Cas12a expression plasmids were transformed into *Escherichia coli* BL21 (DE3) and stored as glycerol stocks. Starter cultures were grown in LB broth with 50µg/ml kanamycin at 37°C for 16h and diluted 1:50 into 400ml LB-kanamycin broth until an OD_600_ of 0.4-0.6 was reached. Cultures were then induced with 1mM isopropyl β-D-1-thiogalactopyranoside (IPTG) and incubated at 25°C for another 16h. Subsequently, cells were harvested by centrifugation at 3,220g for 20min and resuspended in lysis buffer [50mM HEPES, 500mM NaCl, 2mM MgCl_2_, 20mM imidazole, 1% Triton X-100, 1mM DTT, 0.005mg/ml lysozyme (Vivantis), 1X Halt Protease Inhibitor Cocktail (Thermo Fisher Scientific)], followed by sonication at high power for 10 cycles of 30s ON/OFF (Bioruptor Plus; Diagenode). Lysates were clarified by centrifugation at 10,000g for 15min. The supernatants were pooled, loaded onto a gravity flow column packed with Ni-NTA agarose (Qiagen), and rotated for 2h at 4°C. The column was washed twice with 5ml wash buffer (50mM Tris, 300mM NaCl and 30mM imidazole). Five elutions were performed with 500µl elution buffer (50mM Tris, 300mM NaCl, and 200mM imidazole) and analysed by SDS-PAGE. The final gel filtration step was performed with a HiLoad 16/600 Superdex 200pg column (GE Healthcare) on a fast protein liquid chromatography purification system (ÄKTA Explorer; GE Healthcare), which was eluted with storage buffer (50mM Tris, 300mM NaCl, and 1mM DTT). Fractions containing Cas12a were collected, analysed by SDS-PAGE, and concentrated to around 500µl with Vivaspin 20, 50,000 MWCO concentrator units (Sartorius). Glycerol was added to a final concentration of 20%. Protein concentrations were measured with the Quick Start Bradford Protein Assay (Bio-Rad) aliquoted, and stored at -80°C. For comparison, EnGen Lba Cas12a was purchased from New England Biolabs (NEB).

### gRNA design

Complete genomes of SARS-CoV-2 (accession MN908947.3), SARS-CoV (accession NC_004718.3), and MERS-CoV (accession NC_019843.3) were retrieved from NCBI (https://www.ncbi.nlm.nih.gov/) and aligned with MUSCLE (https://www.ebi.ac.uk/Tools/msa/muscle/) using default settings. Potential target sites (20nt spacers) in the ORF1AB, S, and N genes were selected from non-conserved regions containing a TTTV PAM. Potential targets were filtered after a specificity check on BLASTn (https://blast.ncbi.nlm.nih.gov/Blast.cgi) to remove non-specific candidates. Truncated gRNAs were generated by shortening their spacers to 18nt and 19nt lengths at the 3’ end.

### *In vitro* transcription (IVT) of gRNAs

Templates for gRNA synthesis were designed with the following sequence order: T7 promoter-Cas12a scaffold-spacer. Top strand DNA oligos consisting of the T7 promoter (5’-TAATACGACTCACTATAGG-3’) and scaffold (5’-TAATTTCTACTCTTGTAGAT-3’ for AsCas12a and its variants; 5’-AATTTCTACTAAGTGTAGAT-3’ for LbCas12a) were annealed to the bottom strand and extended by Q5 High-Fidelity DNA polymerase (NEB). IVT of the dsDNA products was performed with the HiScribe T7 Quick High Yield RNA Synthesis kit (NEB) at 37°C overnight. Following DNase I digestion, gRNAs were purified with the RNA Clean & Concentrator-5 kit (ZYMO Research), analysed by 2% TAE-agarose gel electrophoresis to assess RNA integrity, measured with NanoDrop 2000, and stored at - 20°C.

### Synthesis of DNA and RNA templates

Gene fragments (gBlocks) were cloned into pCR-Blunt II-TOPO vector using the Zero Blunt TOPO PCR Cloning kit (Invitrogen) and their sequences were verified by Sanger sequencing. The vectors were used as templates for PCR with Q5 High-Fidelity DNA polymerase (NEB) and the products were gel extracted and purified with the PureNA Biospin Gel Extraction kit (Research Instruments). DNA concentrations were measured using NanoDrop 2000 and all the DNA samples were stored at 4°C. To generate RNA templates for RT-LAMP assays, the forward primers used for PCR were appended with the T7 promoter sequence. After PCR amplification with the gBlock-TOPO vectors as template, IVT was performed as described for gRNA generation.

### Fluorescence trans-cleavage assay

Cas ribonucleoprotein (RNP) complexes were pre-assembled with 65nM AsCas12a/ LbCas12a, 195nM gRNA, and 200nM custom ssDNA fluorophore-quencher (FQ) reporter in reaction buffer (1X NEBuffer 3.1 plus 0.4mM DTT) for 30 minutes at room temperature. Subsequently, the cleavage reaction was initiated by adding 3nM DNA template (approximately 1E11 copies) to a total volume of 50µl and then transferred to a 96-well microplate (Costar). Fluorescence intensities were measured with either the Infinite M1000 Pro (Tecan) or the Spectramax M5 plate reader (Molecular Devices) for 30 minutes at room temperature, with measurement intervals of 5 minutes (λ_ex_: 485 nm; λ_em_: 535 nm).

### RT-LAMP reaction

Synthetic SARS-CoV-2 RNA templates were serially diluted and amplified by RT-LAMP using the WarmStart LAMP Kit (NEB). LAMP primers were added to a final concentration of 0.2µM for F3 and B3, 1.6µM for FIP and BIP, and 0.8µM for LF and LB. The optimal temperature for RT-LAMP was found to be 65°C. Subsequently, 4μl RT-LAMP products were used as templates for the trans-cleavage assay, instead of 3nM PCR-amplified DNA template.

### Lateral flow readout

500nM of custom ssDNA biotin reporter was used instead of the FQ reporter. The Cas12a detection reaction was performed at 37°C for 10 minutes. Subsequently, 50μL HybriDetect assay buffer (Milenia Biotec) was added to the reaction and a HybriDetect (Milenia Biotec) dipstick was inserted directly into the solution in an upright position. The dipstick was incubated in the reaction for 2 minutes at room temperature before inspection.

## Acknowledgements

This work is supported by an Industry Alignment Fund – Pre-positioning Programme grant from the Agency for Science, Technology and Research (H18/01/a0/019 to MHT), a Ministry of Education Academic Research Fund Tier 1 grant (2017-T1-001-214 to MHT), an Open Fund - Individual Research Grant from the National Medical Research Council (NMRC/OFIRG/0017/2016 to MHT), and core funding from the Genome Institute of Singapore (to MHT). The authors also thank Xin Ning Nicole Tang for help with setting up the CRISPR diagnostics workflow and members of the DaRE lab for discussions.

## Authors’ contributions

M.H.T conceived the project and provided overall supervision. K.H.O., J.W.D.T., S.Y.T., and M.H.T. designed the experiments. K.H.O., J.W.D.T., S.Y.T., M.M.L., and P.K. performed the experiments. S.J. and Y.-G.G. assisted with protein purification. K.H.O., J.W.D.T., S.Y.T., and M.H.T. analysed the data. M.H.T. wrote the manuscript with inputs from K.H.O., J.W.D.T., and S.Y.T. All authors approved the manuscript.

## Additional information

## Supplementary Information

accompanies this paper

