## Supplementary Information for "A CRISPR-based SARS-CoV-2 diagnostic assay that is robust against viral evolution and RNA editing"

### SUPPLEMENTARY FIGURES

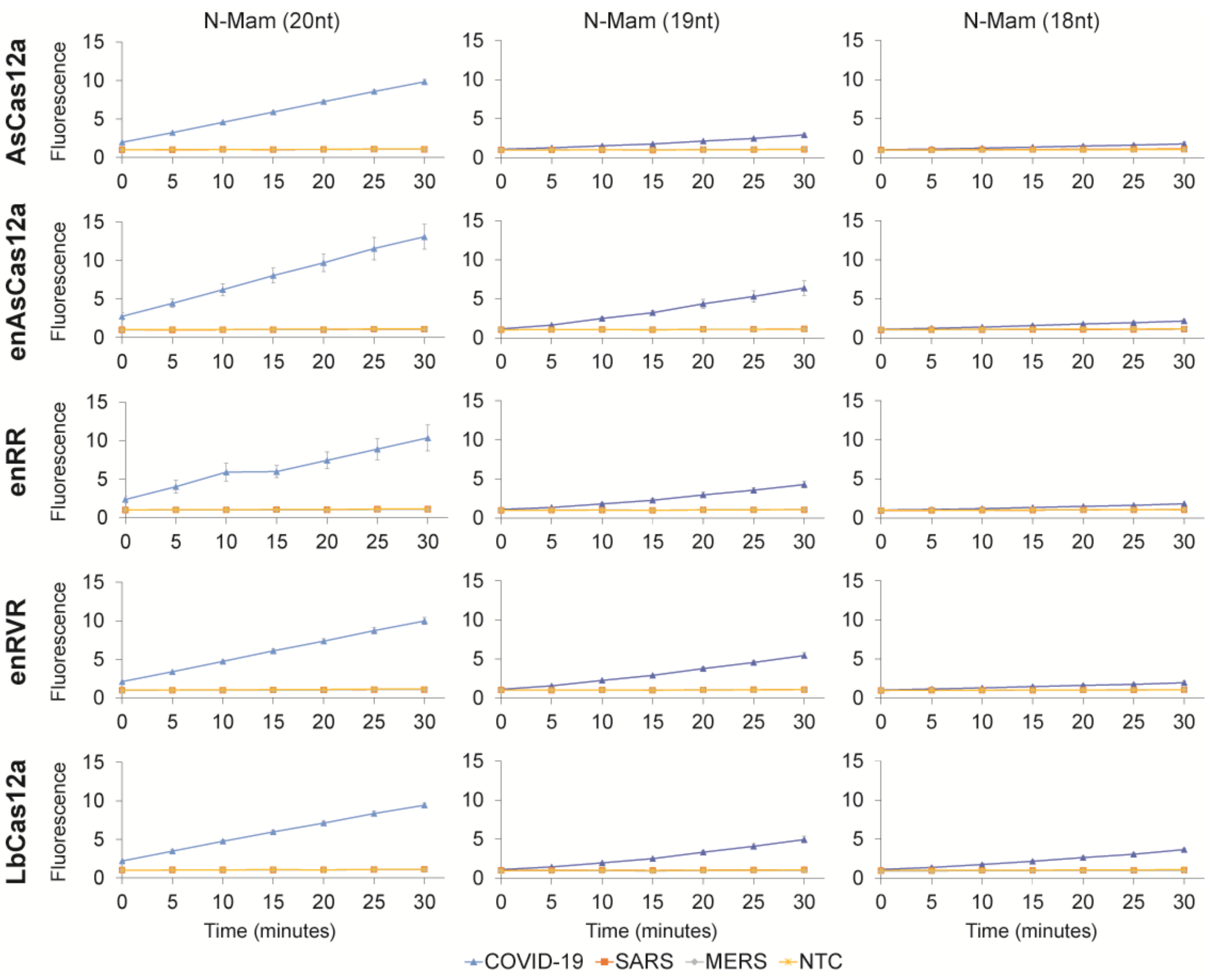


**Supplementary Fig. 1** Time courses of the fluorescence intensity in our trans-cleavage assays for various Cas12a nucleases complexed with perfect matched (PM) N-Mam gRNAs of three different spacer lengths. The assays were performed at 24$^{o}$C and approximately 1E11 copies of purified DNA template were used as input. All the measurements were normalized to the no-template control (NTC) at the start of the experiment. Data represent mean ± s.e.m. (n = 3 biological replicates).

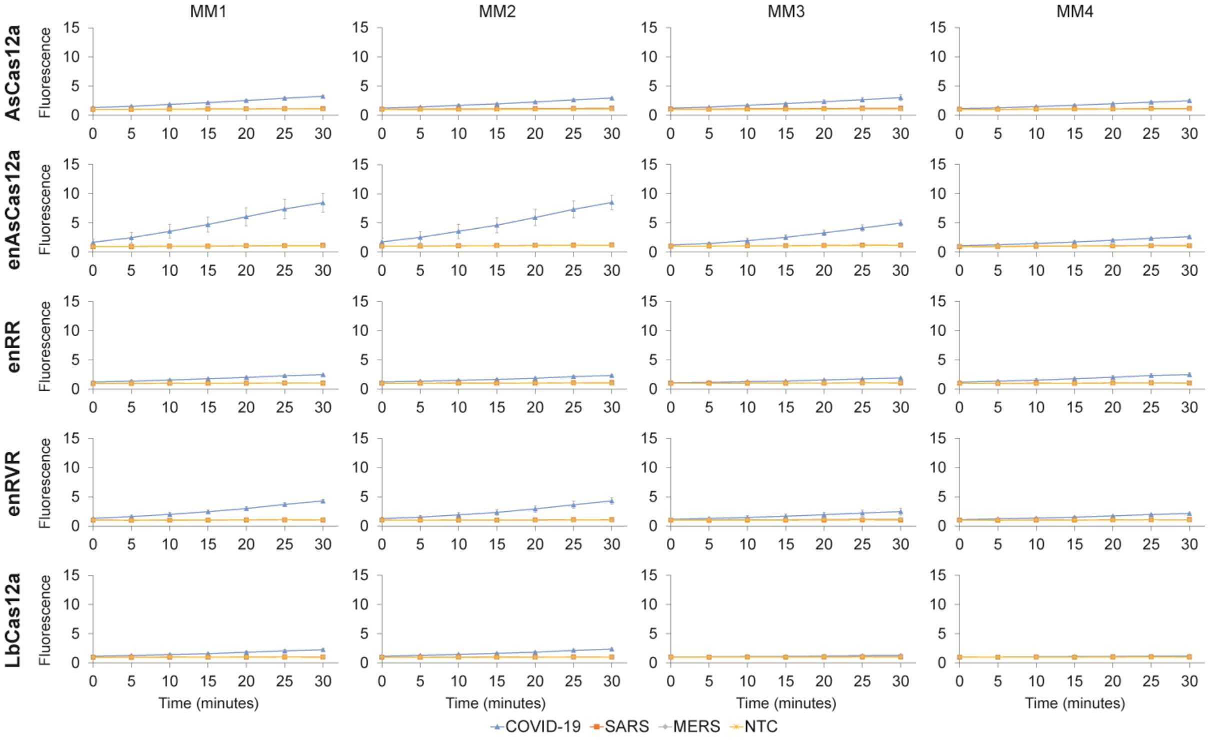
MM1
MM2
MM3
MM4
AsCas12a
enAsCas12a
enRR
enRVR
LbCas12a
Fluorescence
Time (minutes)
COVID-19
SARS
MERS
NTC

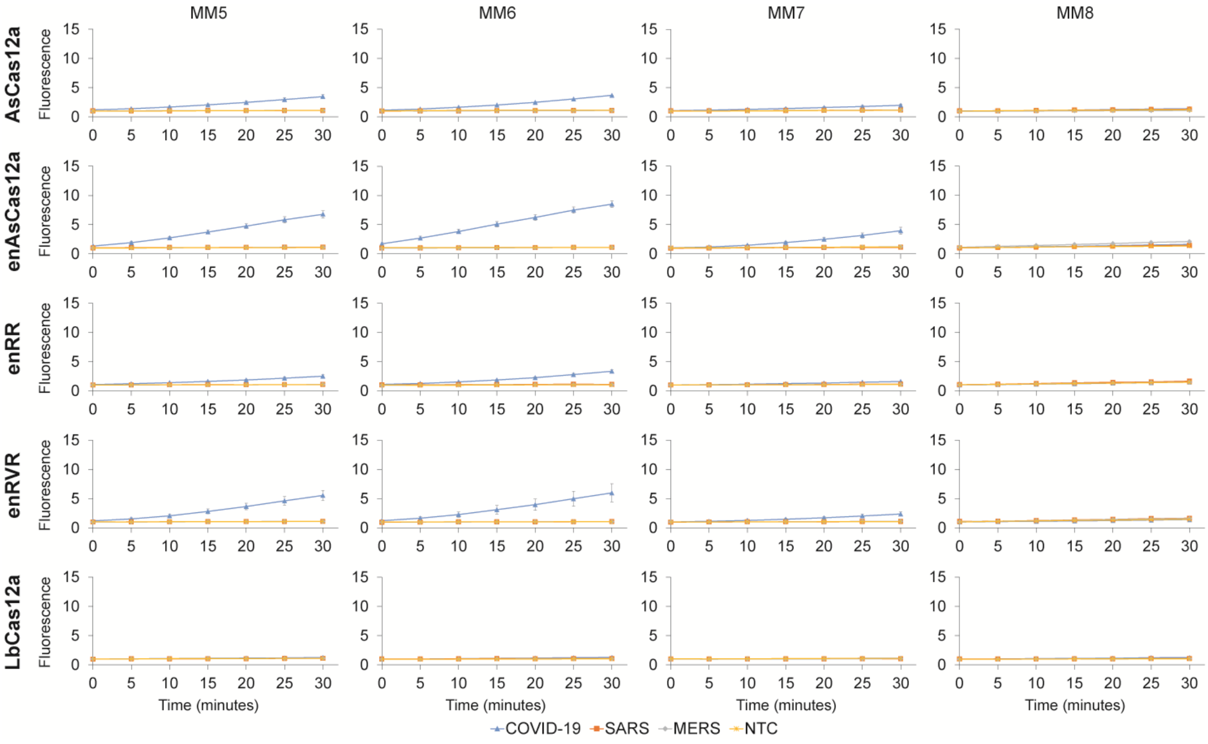
MM5
MM6
MM7
MM8
AsCas12a
enAsCas12a
enRR
enRVR
LbCas12a
Fluorescence
Time (minutes)
COVID-19
SARS
MERS
NTC

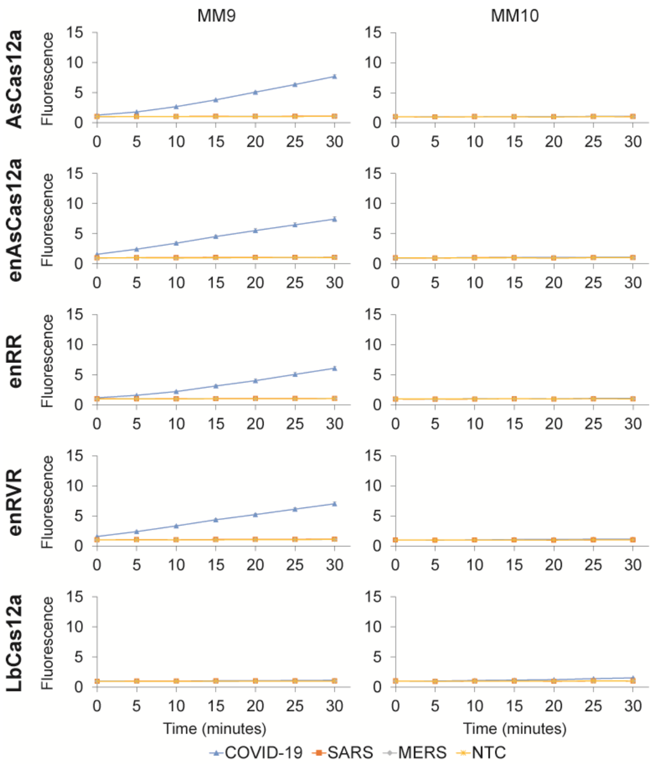


**Supplementary Fig. 2** Time courses of the fluorescence intensity in our trans-cleavage assays for various Cas12a nucleases complexed with mismatched (MM) N-Mam gRNAs of spacer length 20nt. The assays were performed at 24$^{o}$C and approximately 1E11 copies of purified DNA template were used as input. Data represent mean ± s.e.m. (n = 3-4 biological replicates).

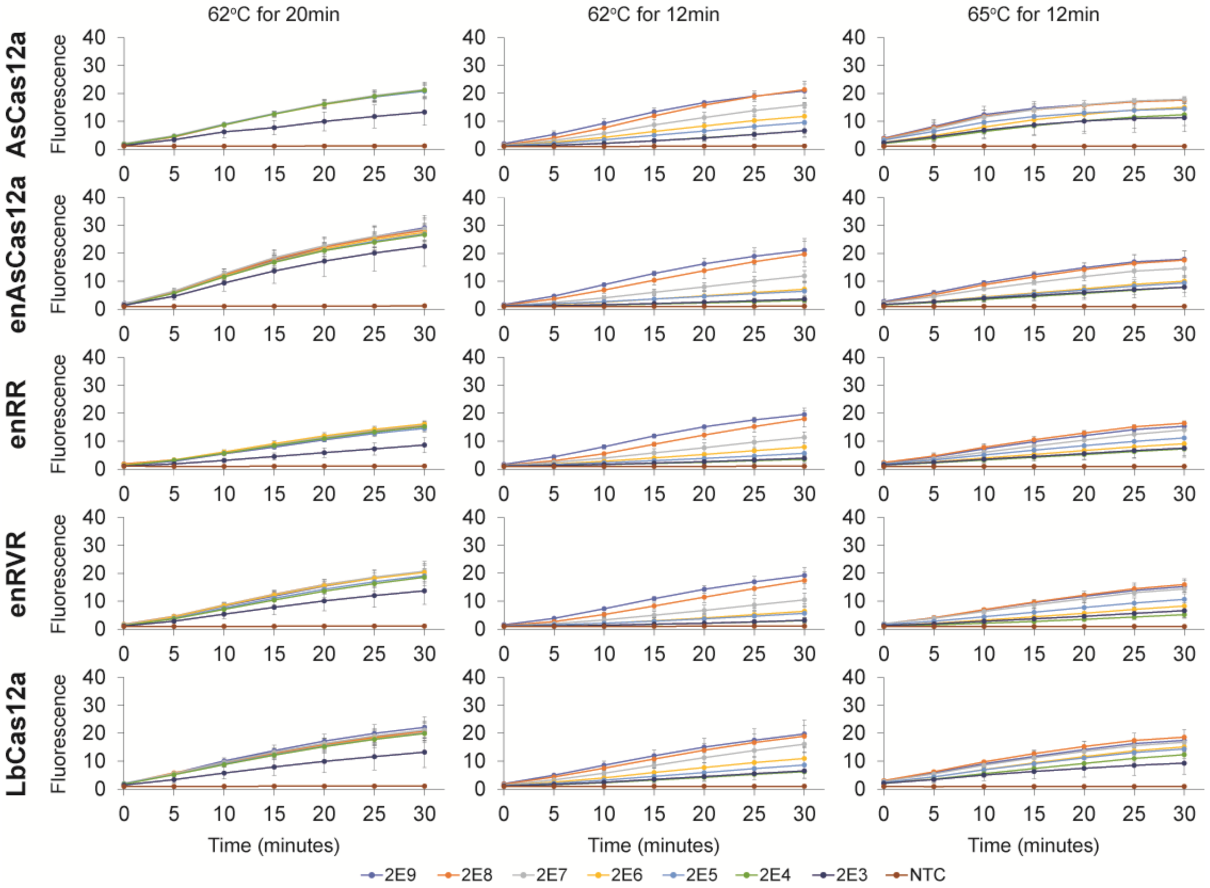


**Supplementary Fig. 3** Time courses of the fluorescence intensity in our trans-cleavage assays for various Cas12a nucleases complexed with perfect matched (PM) N-Mam gRNAs of spacer length 20nt. Before the Cas detection reaction, various copies of *in vitro* transcribed SARS-CoV-2 RNA fragments (see legend) were used as input to an RT-LAMP reaction performed under three different conditions (62$^{o}$C for 20 minutes, 62$^{o}$C for 12 minutes, and 65$^{o}$C for 12 minutes). Subsequently, 4µl LAMP products (out of 25µl) were used for the cleavage assays, which were performed at 24$^{o}$C. Data represent mean ± s.e.m. (62$^{o}$C for 20min: n = 4-7 biological replicates; 62$^{o}$C/ 65$^{o}$C for 12min: n = 3 biological replicates).

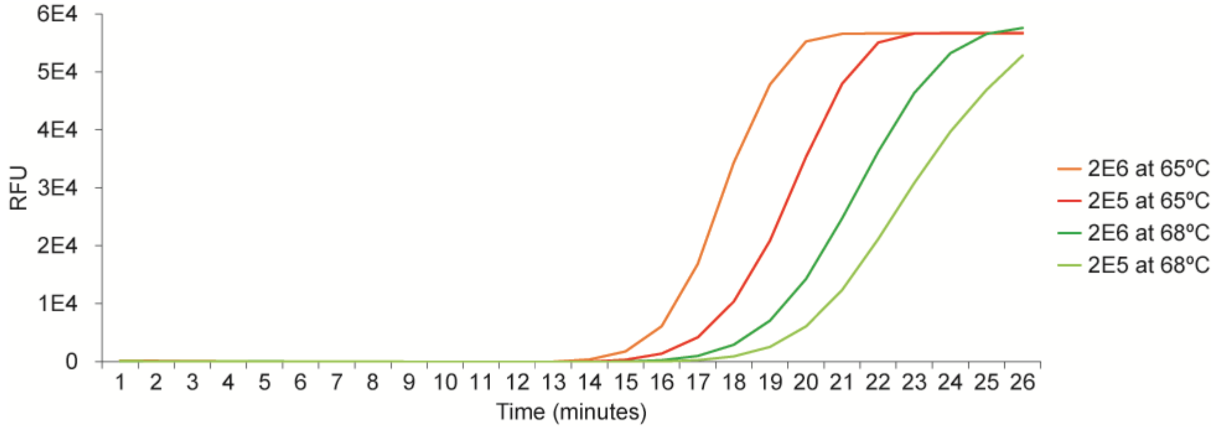


**Supplementary Fig. 4** Real-time monitoring of the RT-LAMP reaction performed at two different temperatures, 65ºC and 68ºC. Fluorescence signal was generated by the addition of a dye that was provided with the WarmStart LAMP kit (New England Biolabs). 2E5 and 2E6 copies of synthetic SARS-CoV-2 RNA input were tested.

**O1 gRNA** (19,968-19,987nt)

| | |
|---|---|
| SARS-CoV-2 (MN908947.3) | **TTTG**TGCACCACTCACTGTCTTTT |
| SARS-CoV (NC_004718.3) | CTTGTTCTTCACTTACTGTCTTGT |
| MERS-CoV (NC_019843.3) | TATGT------------------T |

**O2 gRNA** (20,074-20,093nt)

| | |
|---|---|
| SARS-CoV-2 (MN908947.3) | **TTTA**CAACCAT----CTGTAGGTCCCAA |
| SARS-CoV (NC_004718.3) | TCTAACACCTT----CAAAGGGACCAGC |
| MERS-CoV (NC_019843.3) | ---AATACCCTTGTATGGTAGGTCCTGA |

**S1 gRNA** (22,286-22,305nt)

| | |
|---|---|
| SARS-CoV-2 (MN908947.3) | **TTTA**CTTGCTTTACATAGAA---------GTTA |
| SARS-CoV (NC_004718.3) | C-------------------------------- |
| MERS-CoV (NC_019843.3) | CTTGCCTGTTTATGATACTATTAAGTATTATTC |

**S2 gRNA** (22,310-22,329nt)

| | |
|---|---|
| SARS-CoV-2 (MN908947.3) | **TTTG**ACTCCTGGTGATTCTTCTTC |
| SARS-CoV (NC_004718.3) | ----ATTCTTACAGCCTTTTCACC |
| MERS-CoV (NC_019843.3) | TATCATTCCTCACAGTATTCGTTC |

**S3 gRNA** (22,518-22,537nt)

| | |
|---|---|
| SARS-CoV-2 (MN908947.3) | **TTTA**GAGTCCAACCAACAGAATCT |
| SARS-CoV (NC_004718.3) | TTCAGGGTTGTTCCCTCAGGAGAT |
| MERS-CoV (NC_019843.3) | TTCGAAGCAAAACCTTCTGGCTCA |

**N-Mam gRNA** (29,196-29,215nt)

| | |
|---|---|
| SARS-CoV-2 (MN908947.3) | **TTTG**CCCCCAGCGCTTCAGCGTTC |
| SARS-CoV (NC_004718.3) | TTTGCTCCAAGTGCCTCTGCATTC |
| MERS-CoV (NC_019843.3) | CTTGCTCCTACAGCCAGTGCTTTT |

**N1 gRNA** (29,312-29,331nt)

| | |
|---|---|
| SARS-CoV-2 (MN908947.3) | **TTTC**AAAGATCAAGTCATTTTGCT |
| SARS-CoV (NC_004718.3) | ATTCAAAGACAACGTCATACTGCT |
| MERS-CoV (NC_019843.3) | CTACAATAAGTGGTTGGAGCTTCT |

**Supplementary Fig. 5** Multiple sequence alignment of the various gRNAs evaluated in our study across SARS-CoV-2, SARS-CoV, and MERS-CoV. The TTTV PAM is indicated in bold blue before each 20nt spacer. The mismatches are also shown in red.

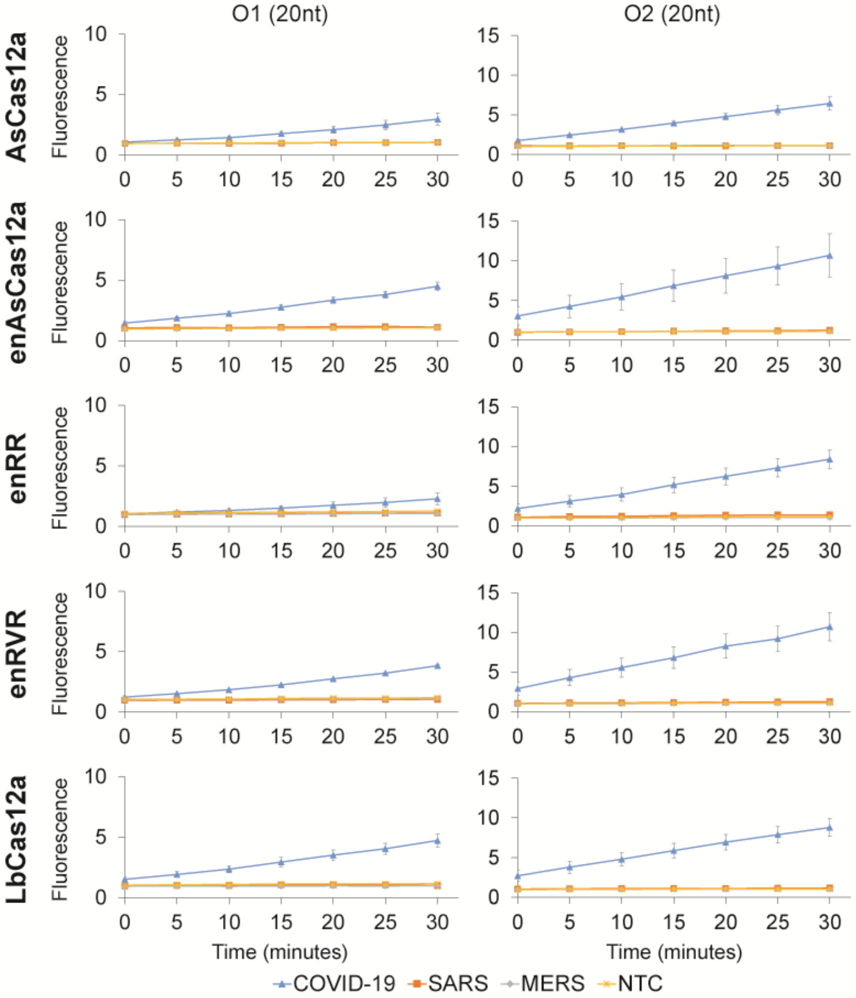

O1 (20nt)
O2 (20nt)
AsCas12a
enAsCas12a
enRR
enRVR
LbCas12a
Fluorescence
Time (minutes)
COVID-19
SARS
MERS
NTC

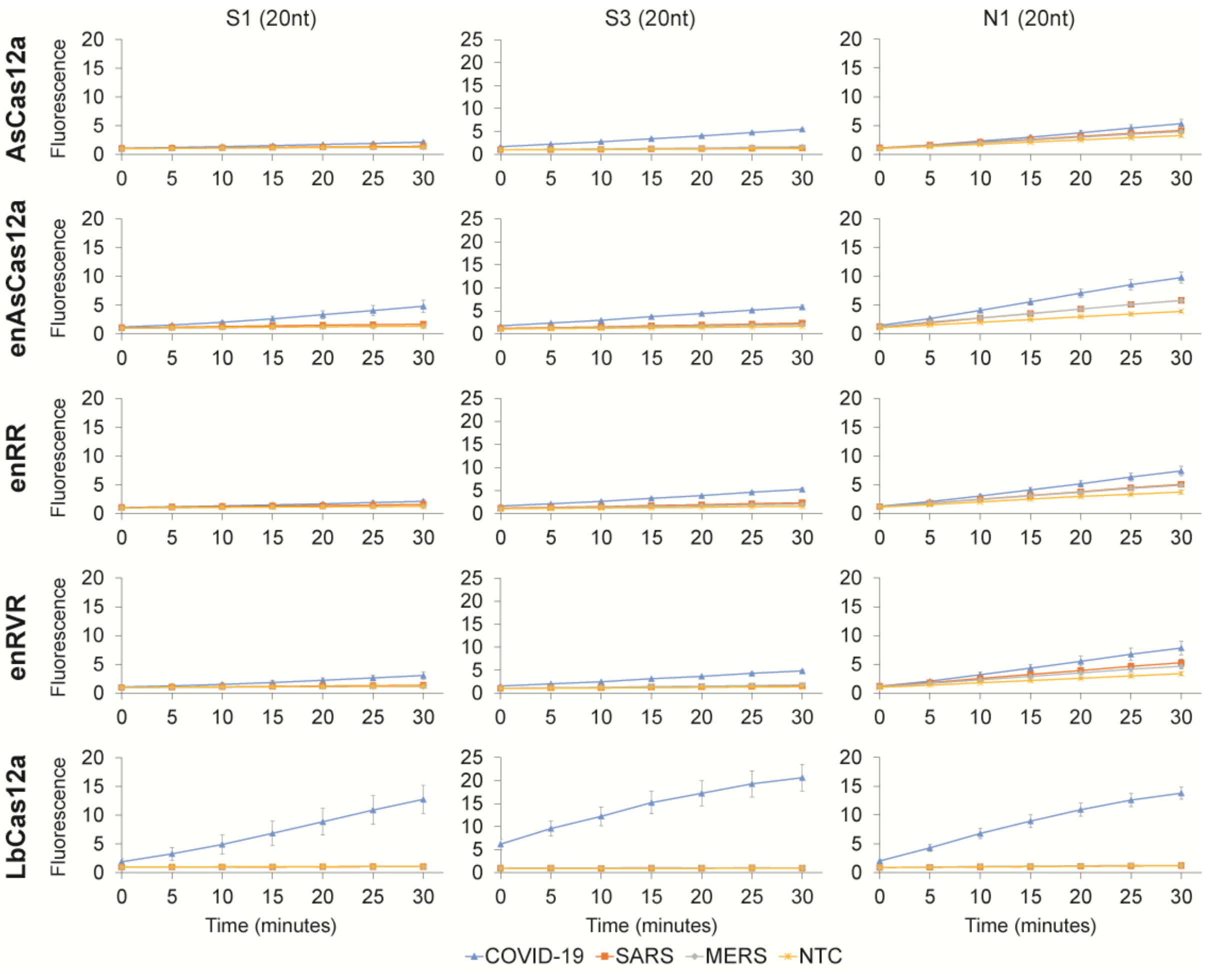


**Supplementary Fig. 6** Time courses of the fluorescence intensity in our trans-cleavage assays for various Cas12a nucleases complexed with perfect matched (PM) gRNAs targeting the O1, O2, S1, S3, and N1 loci of the SARS-CoV-2 genome. The assays were performed at 24ºC and approximately 1E11 copies of purified DNA template were used as input. All the measurements were normalized to the no-template control (NTC) at the start of the experiment. Data represent mean ± s.e.m. (n = 3-6 biological replicates).

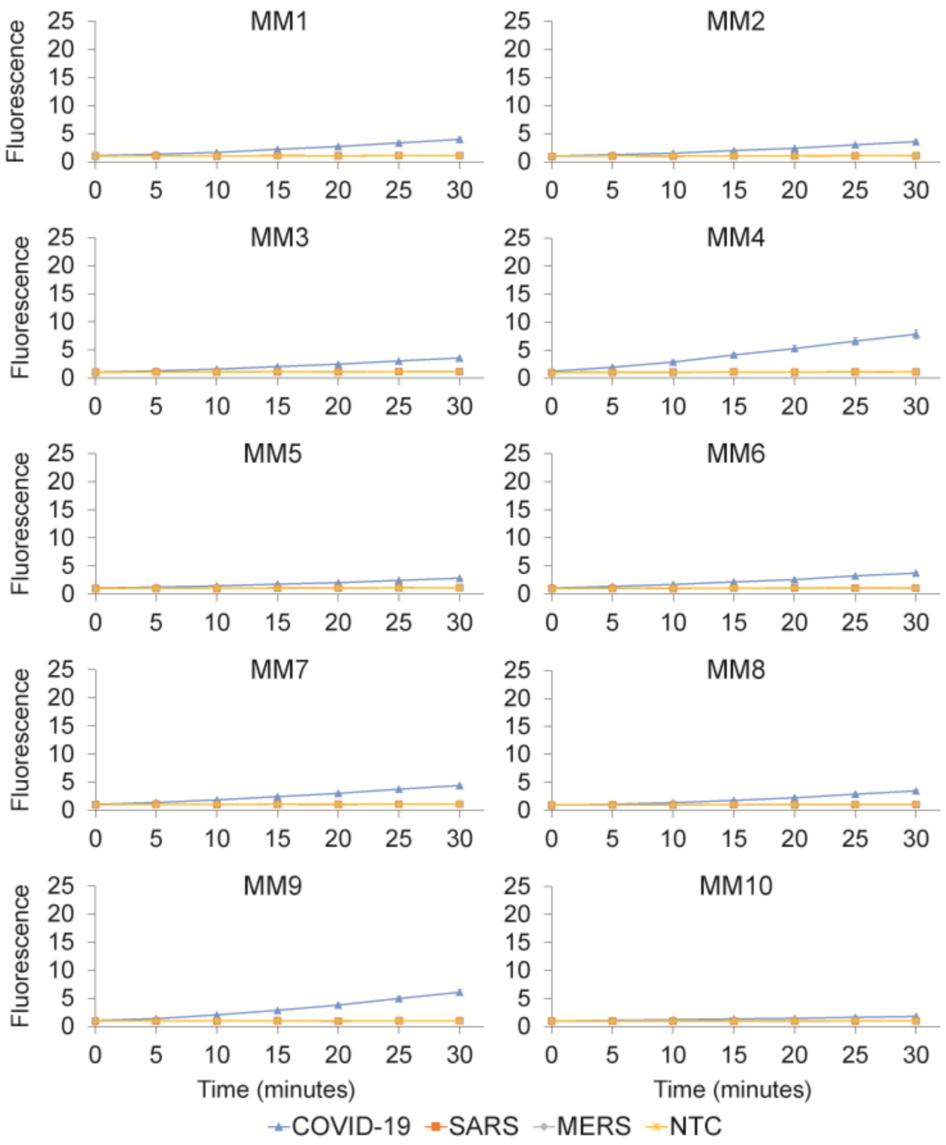


**Supplementary Fig. 7** Time courses of the fluorescence intensity in our trans-cleavage assays for various Cas12a nucleases complexed with mismatched (MM) S3 gRNAs of spacer length 20nt. The assays were performed at 24°C and approximately 1E11 copies of purified DNA template were used as input. Data represent mean ± s.e.m. (n = 3 biological replicates).

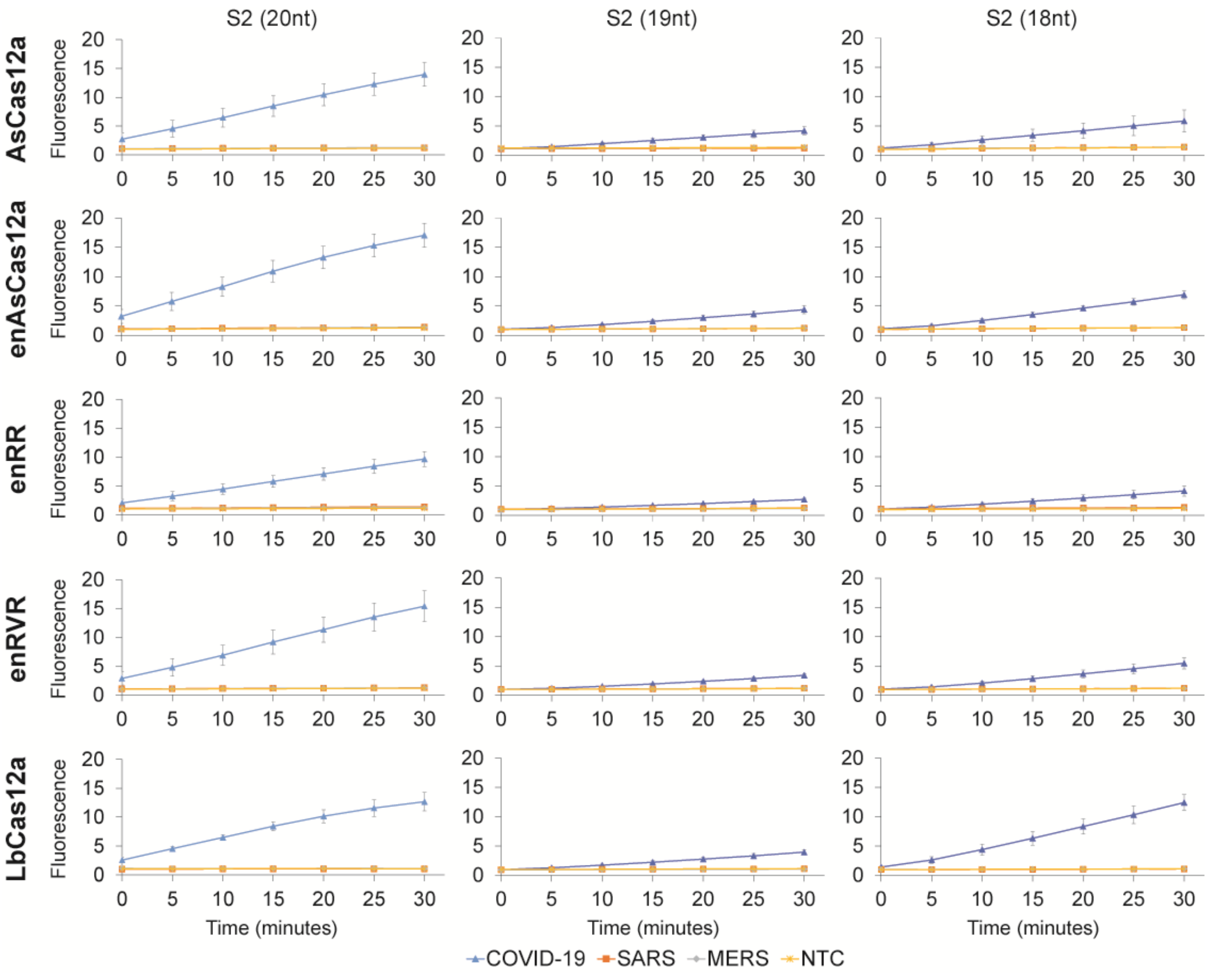


**Supplementary Fig. 8** Time courses of the fluorescence intensity in our trans-cleavage assays for various Cas12a nucleases complexed with perfect matched (PM) S2 gRNAs of three different spacer lengths. The assays were performed at 24°C and approximately 1E11 copies of purified DNA template were used as input. All the measurements were normalized to the no-template control (NTC) at the 0min timepoint. Data represent mean ± s.e.m. (n = 3-4 biological replicates).

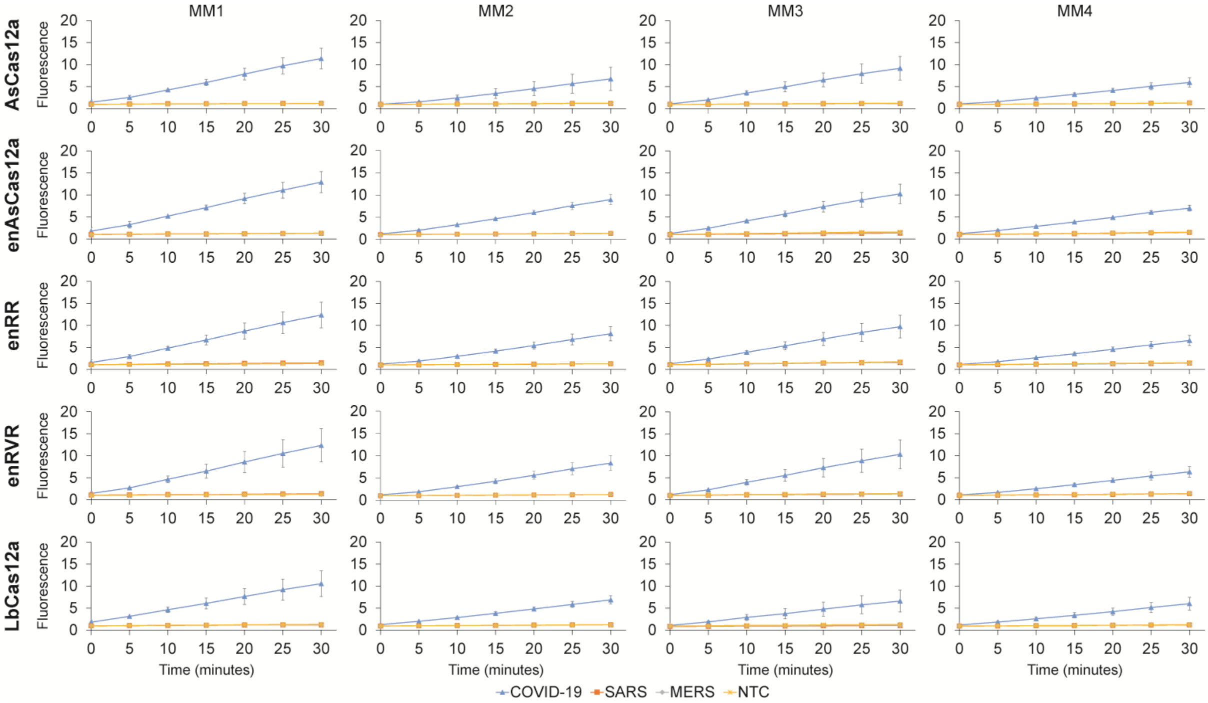

MM1
MM2
MM3
MM4
AsCas12a
enAsCas12a
enRR
enRVR
LbCas12a
Fluorescence
Time (minutes)
COVID-19
SARS
MERS
NTC


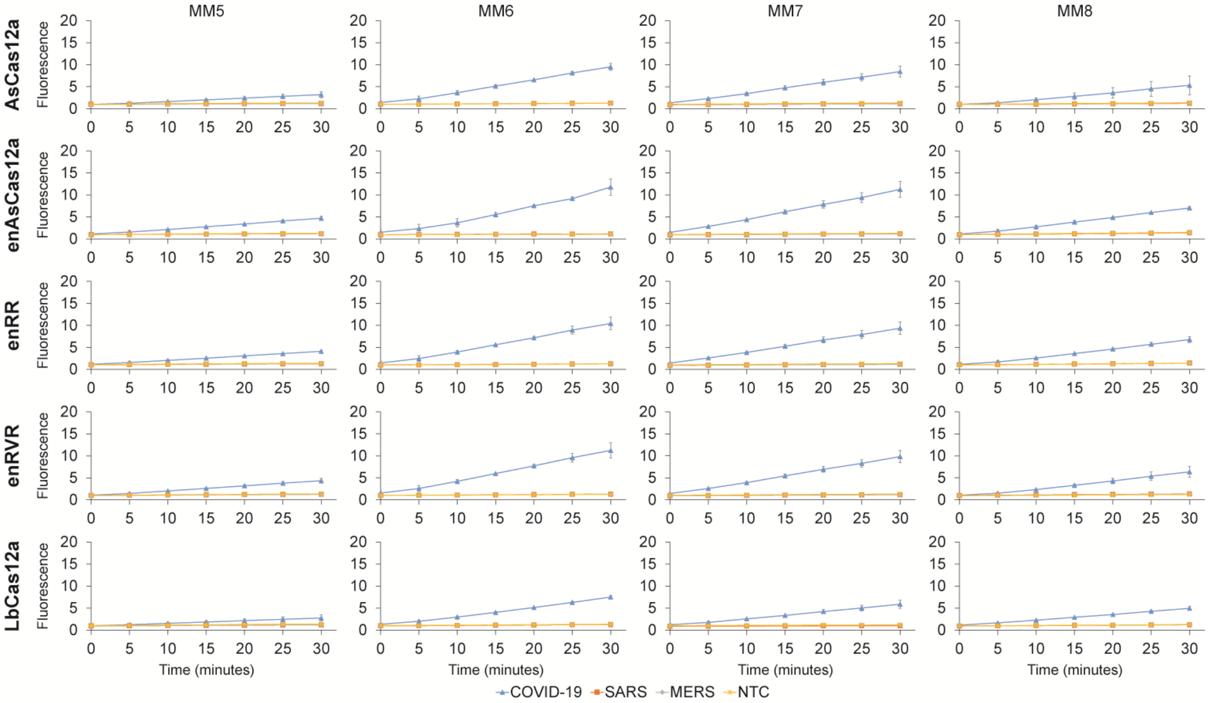

MM5
MM6
MM7
MM8
AsCas12a
enAsCas12a
enRR
enRVR
LbCas12a
Fluorescence
Time (minutes)
COVID-19
SARS
MERS
NTC

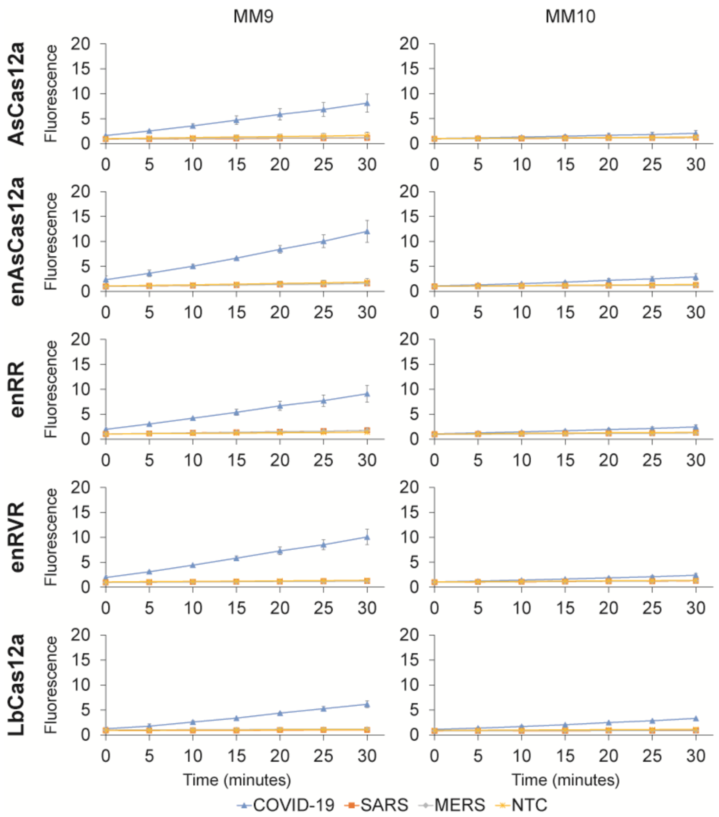


**Supplementary Fig. 9** Time courses of the fluorescence intensity in our trans-cleavage assays for various Cas12a nucleases complexed with mismatched (MM) S2 gRNAs of spacer length 20nt. The assays were performed at 24ºC and approximately 1E11 copies of purified DNA template were used as input. Data represent mean ± s.e.m. (n = 2 biological replicates).

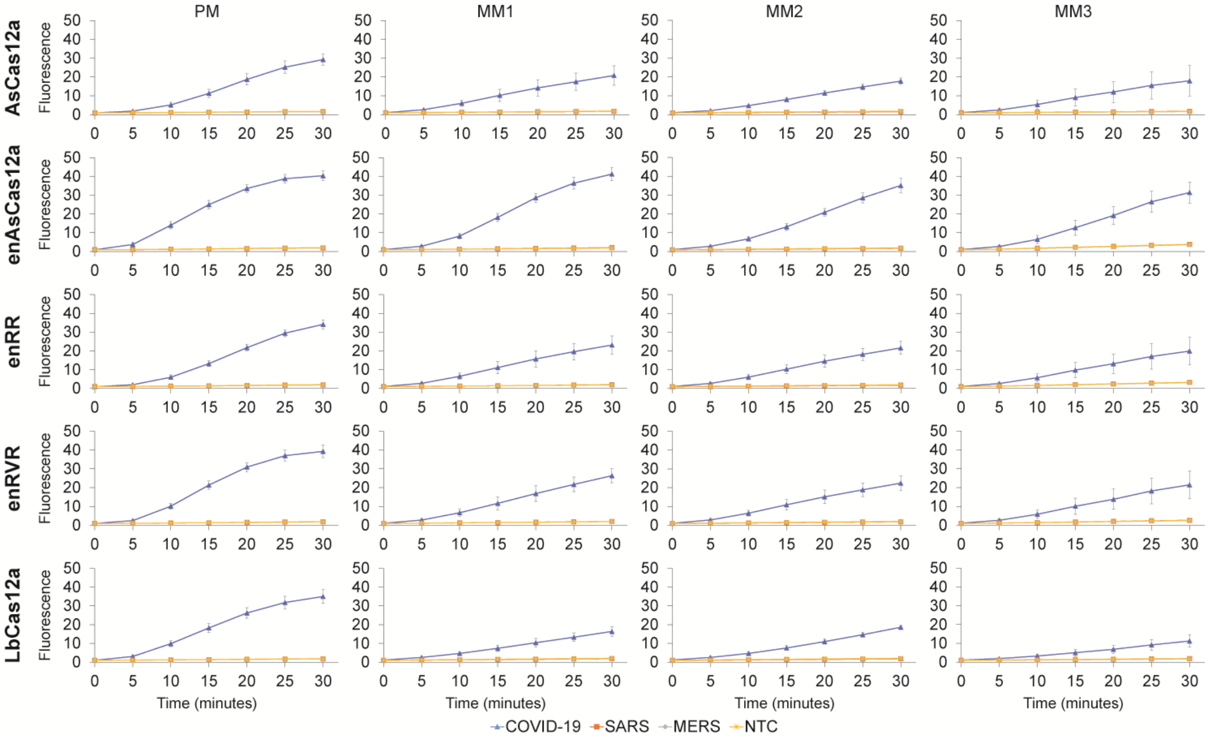
PM
MM1
MM2
MM3
AsCas12a
enAsCas12a
enRR
enRVR
LbCas12a
Fluorescence
Time (minutes)
COVID-19
SARS
MERS
NTC

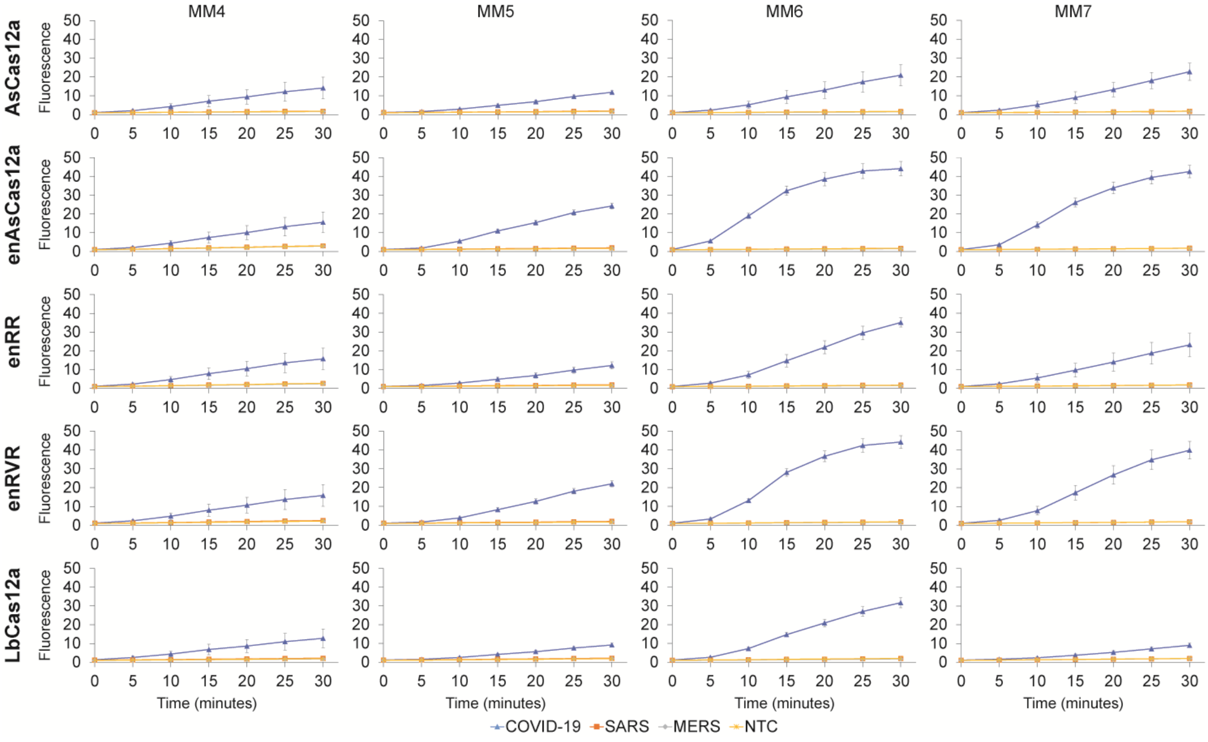
MM4
MM5
MM6
MM7
AsCas12a
enAsCas12a
enRR
enRVR
LbCas12a
Fluorescence
Time (minutes)
COVID-19
SARS
MERS
NTC

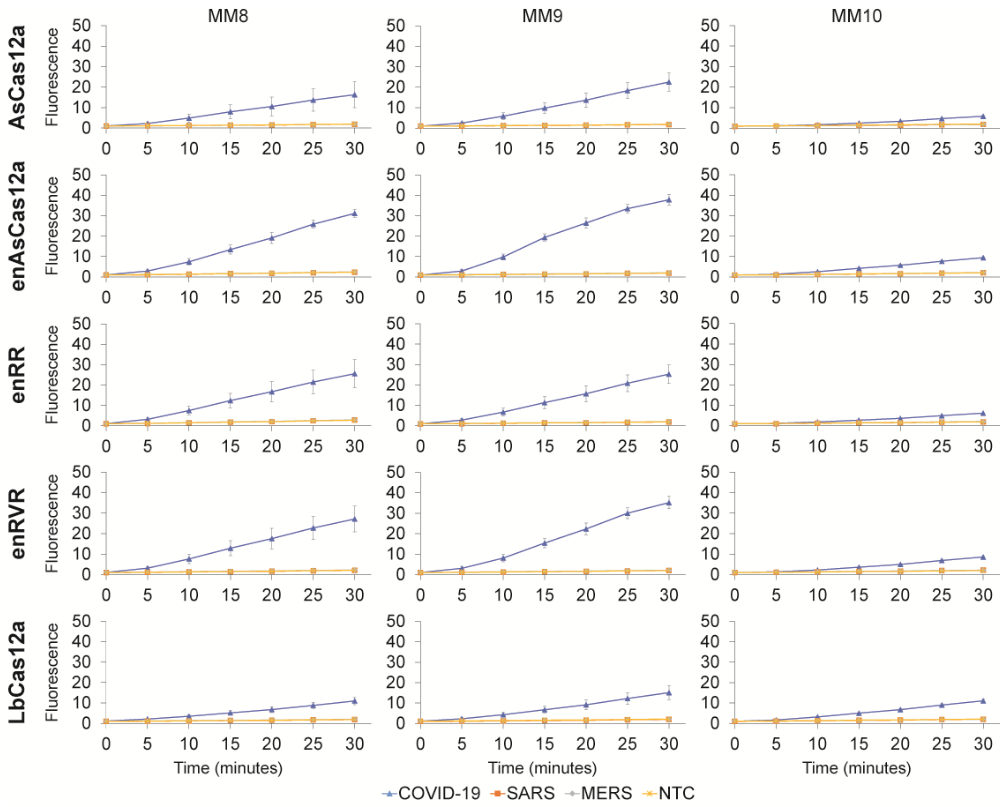


**Supplementary Fig. 10** Time courses of the fluorescence intensity in our trans-cleavage assays for various Cas12a nucleases complexed with either perfect matched (PM) or mismatched (MM) S2 gRNAs of spacer length 20nt. The assays were performed at 37$^{o}$C and approximately 1E11 copies of purified DNA template were used as input. All the measurements were normalized to the no-template control (NTC) at the 0min timepoint. We observed that when the Cas detection reaction was performed at 37$^{o}$C, it proceeded around twice as fast as the same reaction at 24$^{o}$C (room temperature). However, in the event that only one heat block is available (which will have to be used for the RT-LAMP reaction), then the Cas detection reaction can still be carried out at room temperature, although the time has to be lengthened. Data represent mean ± s.e.m. (n = 3 biological replicates).

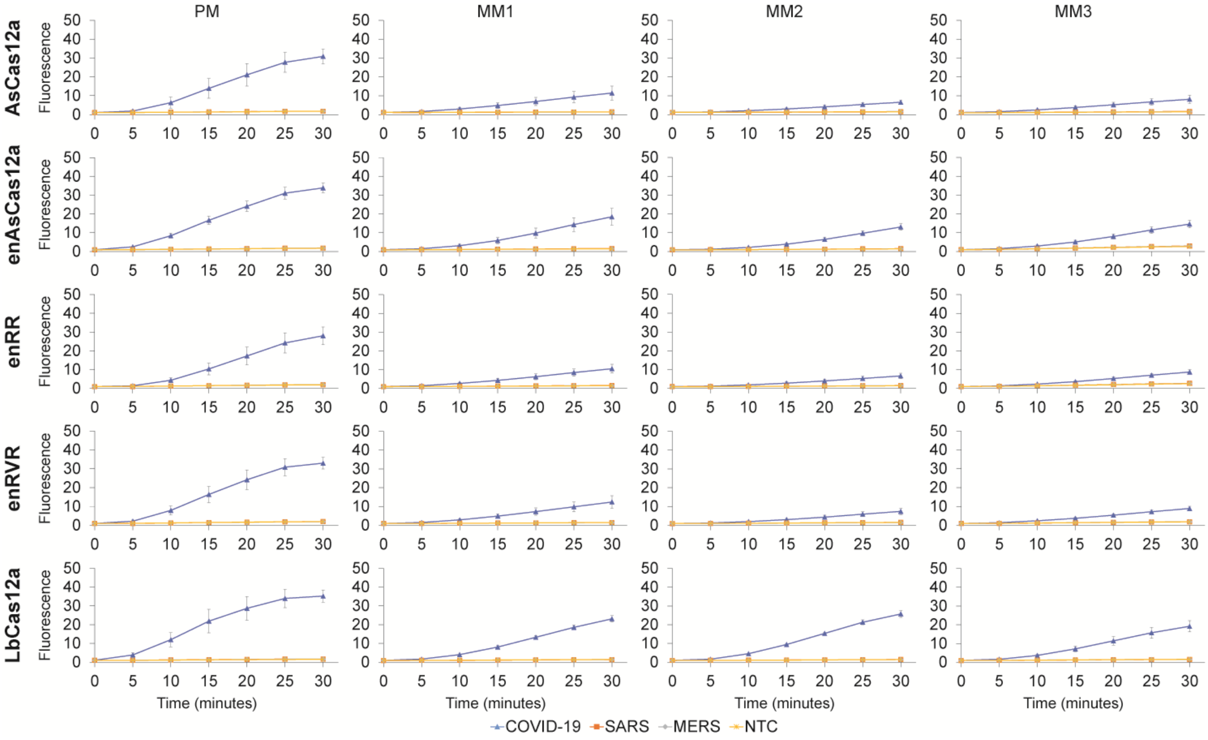
PM
MM1
MM2
MM3
AsCas12a
enAsCas12a
enRR
enRVR
LbCas12a
Fluorescence
Time (minutes)
COVID-19
SARS
MERS
NTC

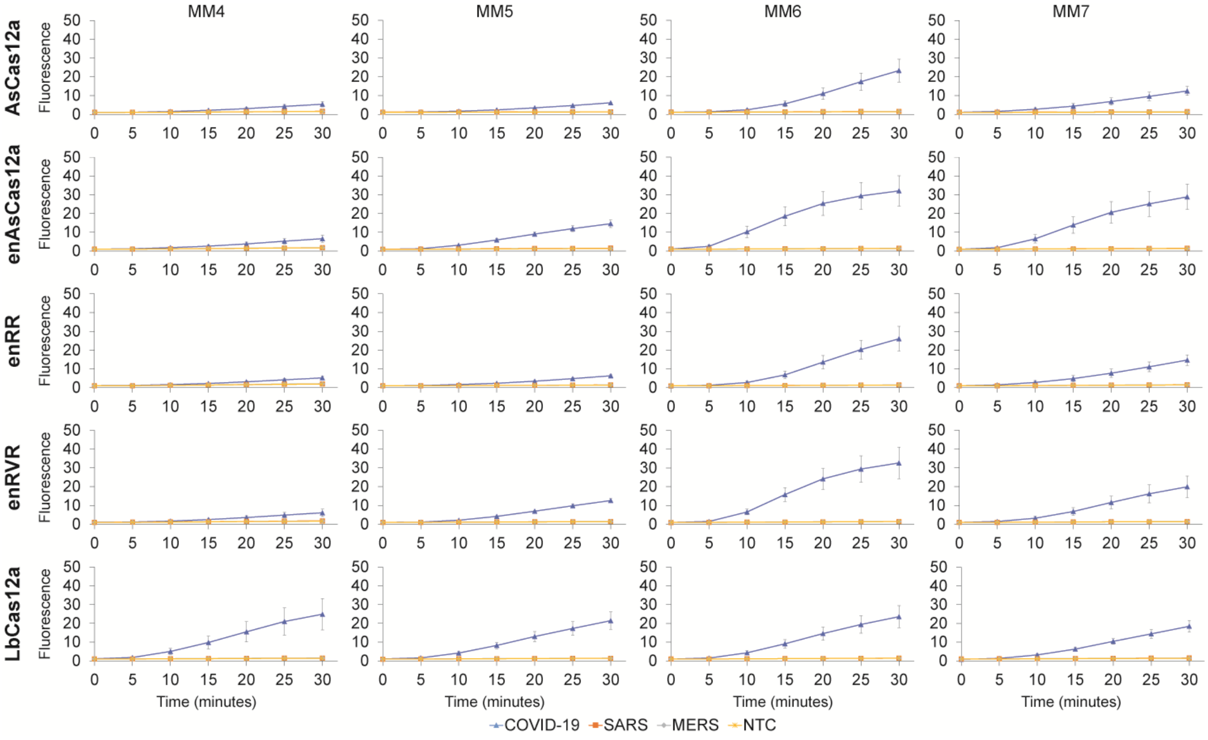
MM4
MM5
MM6
MM7
AsCas12a
enAsCas12a
enRR
enRVR
LbCas12a
Fluorescence
Time (minutes)
COVID-19
SARS
MERS
NTC

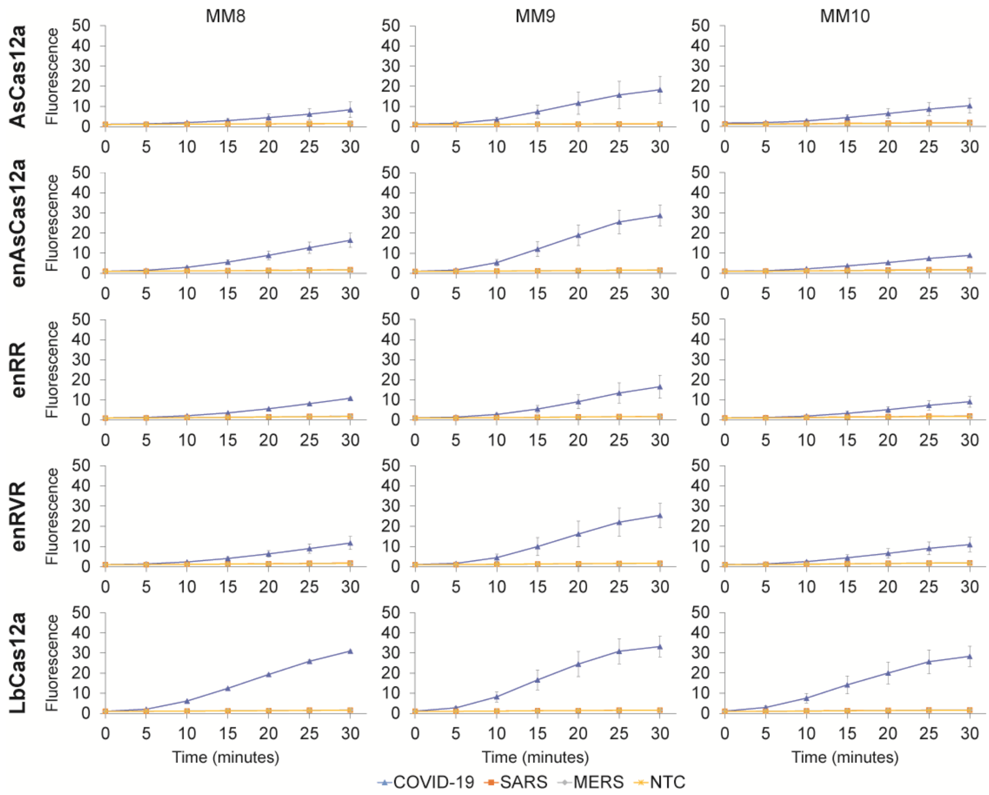


**Supplementary Fig. 11** Time courses of the fluorescence intensity in our trans-cleavage assays for various Cas12a nucleases assembled with two gRNAs – S1 PM gRNA and S2 PM or MM gRNA. All the guides tested here have a spacer length of 20nt. The assays were performed at 37$^{o}$C and approximately 1E11 copies of purified DNA template were used as input. Data represent mean ± s.e.m. (n = 3 biological replicates).

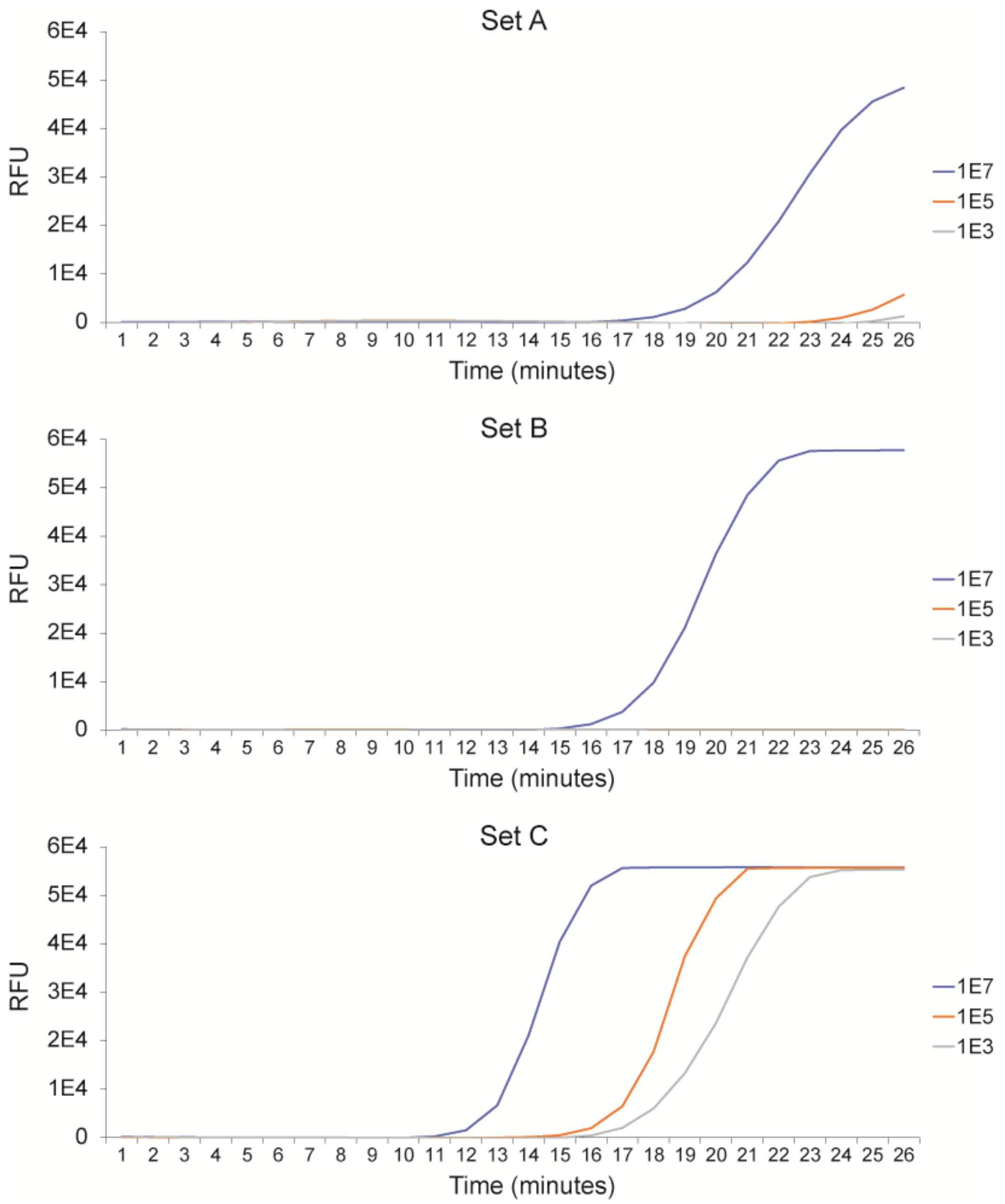


**Supplementary Fig. 12** Real-time monitoring of the RT-LAMP reaction performed at 65$^{o}$C for three different sets of primers targeting the S gene of SARS-CoV-2. Fluorescence signal was generated by the addition of a dye that was provided with the WarmStart LAMP kit (New England Biolabs). 1E3, 1E5, and 1E7 copies of synthetic SARS-CoV-2 RNA input were tested.

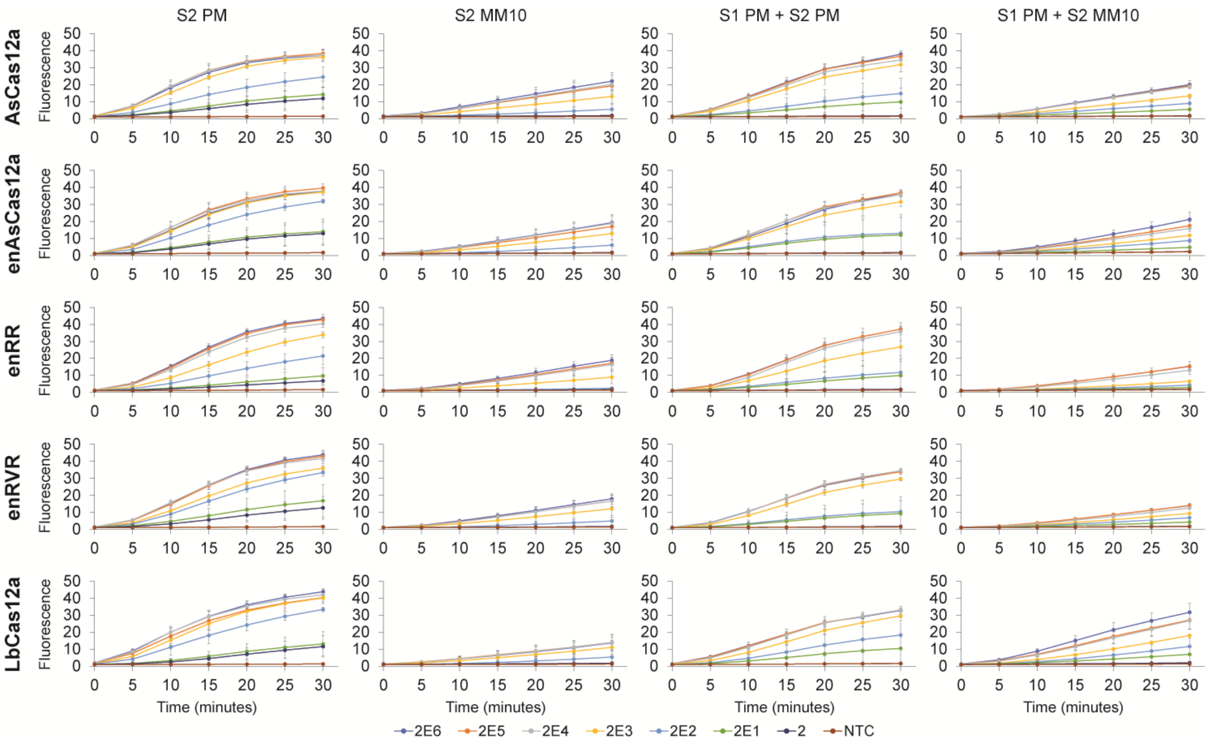


**Supplementary Fig. 13** Time courses of the fluorescence intensity in our trans-cleavage assays for various Cas12a nucleases assembled with S2 PM gRNA alone, S2 MM10 gRNA alone, S1 PM gRNA and S2 PM gRNA, and S1 PM gRNA and S2 MM10 gRNA. All the guides tested here have a spacer length of 20nt. Before the Cas detection reaction, various copies of *in vitro* transcribed SARS-CoV-2 RNA fragments (see legend) were used as input to an RT-LAMP reaction performed at 65$^o$C for 15 minutes. Subsequently, 4μl LAMP products (out of 25μl) were used for the cleavage assays, which were performed at 37$^o$C. Data represent mean ± s.e.m. (n = 3-5 biological replicates).

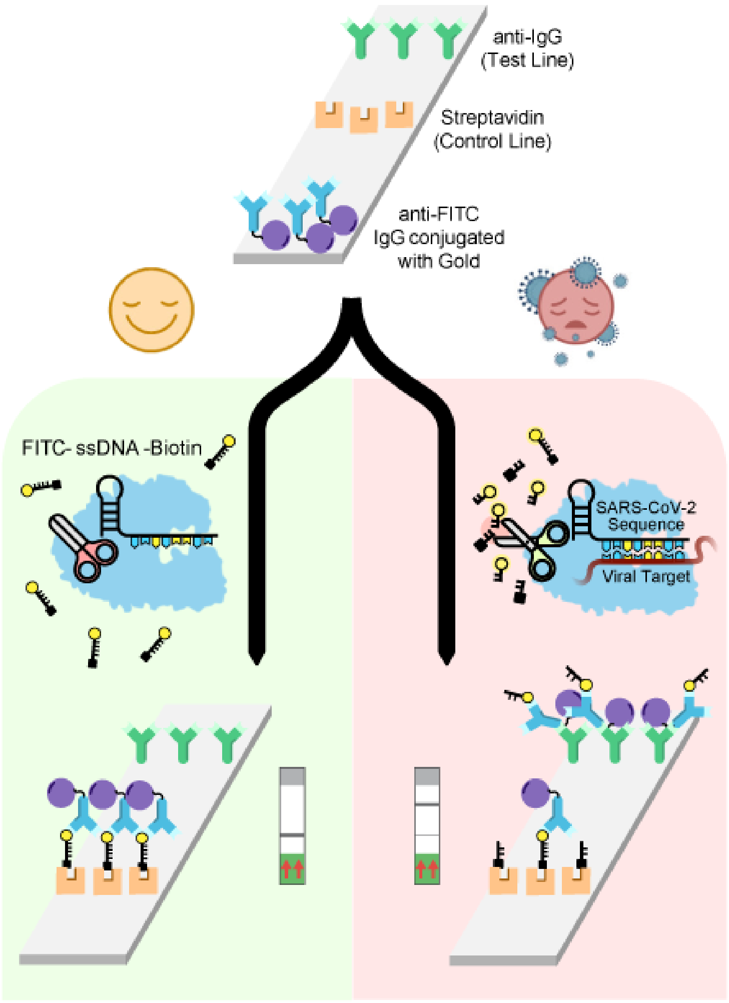


**Supplementary Fig. 14** An illustration of the operating principle of a lateral flow strip or dipstick. On each strip are gold-conjugated IgG antibodies against the fluorophore near the sample pad, streptavidin immobilized at the control line, and antibodies against IgG immobilized at the test line. In the case of a virus-free sample, the Cas nuclease remains inactive and thus the reporter, comprising a fluorophore linked to biotin by a short piece of single-stranded DNA (ssDNA), stays intact. When the reaction is loaded on the strip, the gold-conjugated IgG first binds to the fluorophore and then the entire IgG-reporter complex is captured at the control line due to the high affinity of streptavidin for biotin. Consequently, a dark band is observed at the control line. However, in the case of an infected sample, the Cas nuclease cleaves its viral target, becomes hyperactivated, and then proceeds to cut the linker between the fluorophore and biotin. Subsequently, when the reaction is loaded on the strip, the gold-conjugated IgG still binds to the fluorophore, but the gold will not be deposited at the control line as the fluorophore is now free of biotin. Instead, the IgG-fluorophore complex continues flowing along the strip to the test line, where it is captured by the anti-IgG antibodies there. Consequently, a dark band is observed at the test line.
